# Sustainable Regeneration of 20 Aminoacyl-tRNA Synthetases in a Reconstituted System Toward Self-Synthesizing Artificial Systems

**DOI:** 10.1101/2024.10.03.616507

**Authors:** Katsumi Hagino, Keiko Masuda, Yoshihiro Shimizu, Norikazu Ichihashi

## Abstract

In vitro construction of self-reproducible artificial systems is a major challenge in bottom-up synthetic biology. Here, we developed a reconstituted system capable of sustainably regenerating all 20 aminoacyl-tRNA synthetases (aaRSs), which are major components of the translation system. To achieve this, we needed five types of improvements: 1) optimization of aaRS sequences for efficient translation, 2) optimization of the composition of the translation system to enhance translation, 3) employment of another bacterial AlaRS and SerRS to improve each aminoacylation activity, 4) diminishing the translational inhibition caused by certain aaRS sequences by codon optimization and EF-P addition, and 5) balancing the DNA concentrations of 20 aaRSs to match each requirement. After these improvements, we succeeded in the sustainable regeneration of all 20 aaRSs for up to 20 cycles of 2.5-fold serial dilutions. These methodologies and results provide a substantial advancement toward the realization of self-reproducible artificial systems.

## Introduction

Self-reproduction is a fundamental characteristic of living organisms and its implementation in artificial systems remains a major challenge (*1–17*). Although significant progress has been made in reconstituting biological functions such as DNA replication, metabolism, cellular proliferation, and cytoskeletal function in vitro, the ability to continuously reproduce essential biomolecules such as DNA, RNA, and proteins has not been achieved in synthetic systems (*18–34*). In living organisms, self-replication relies on the synthesis of polymers from small molecules via processes such as DNA replication, transcription, and translation.

To construct a self-reproducible artificial system, the reconstituted transcription/translation system of *Escherichia coli* (PURE system) is a reasonable starting point, because it consists of minimal elements of transcription and translation (*35*). The PURE system consists of macromolecules, including ribosomes, tRNAs, 20 aminoacyl-tRNA synthetases (aaRS), translational initiation/elongation/releasing factors, and T7 RNA polymerase. If all these RNAs and proteins in the PURE system could be sustainably regenerated by the PURE system itself, it would be much closer to a self-reproducible system.

Several studies have successfully synthesized translation proteins (TPs) using the PURE system. Awai *et al.* expressed each of 20 aaRS in the PURE system and detected activity except for PheRS (*36*). Wei and Endy expressed each of 36 TPs in the PURE system and detected each activity of 19 of 23 testable TPs after purification (*37*). Li *et al.* co-expressed most of the ribosomal proteins in one pot and detected reconstitution of the 30S subunit (*38*). Libicher *et al.* co-expressed multiple TPs in the PURE system from three large plasmids encoding most of the Translation factors (*39*), although the expression amounts were insufficient to regenerate the original amount of TPs, and the activities of the expressed TPs were not verified. Doerr *et al.* also reported that a significant portion of the co-expressed proteins was truncated owing to inefficient ribosome processivity (*40*). However, most of these studies did not demonstrate regeneration, because the expressed translation factors were not used for further translation.

As a study demonstrating the regeneration of TPs during transcription/translation reactions, Libicher performed partial regeneration of T7 RNA polymerase and adenylate kinase, as well as 12 aaRSs and RF1, to the second generation by serial dilution experiments using the PURE system in which the target proteins were removed (*41*). In addition, Lavickova *et al.* demonstrated the sustainable self-regeneration of T7 RNA polymerase or up to seven aaRSs using a microfluidic reactor, which allowed continuous expression of GFP for more than 24 h (*42*). Inspired by these studies, we recently constructed a system that couples the expression of 20 aaRSs with replication of DNA encoding aaRSs. Although DNA replication was sustained over four generations in this system, translation activity rapidly declined over generations, indicating that the 20 aaRSs were not sufficiently regenerated (*43*). In summary, sustainable regeneration in PURE systems is still limited to seven aaRSs (*42*), and increasing this number is the next most important issue.

In this study, we attempted to increase the number of regenerative TPs up to all 20 aaRSs, which comprises approximately half of the TPs in the PURE system, except for ribosomes. For this purpose, we first quantified the translation efficiency and specific activity of the aaRSs expressed in the PURE system. Second, we improved the aaRS sequences to increase translation efficiency. Third, we found and employed highly active AlaRS and SerRS from *Geobacillus stearothermophilus*. Fourth, we found that some aaRS sequences had inhibitory effects on the translation activity of the PURE system, and mitigated this effect by removing rare codons and adding EF-P. Fifth, we further optimized the DNA concentration ratio to achieve ideal aaRS expression balance. After these improvements, we succeeded in maintaining the translation activity in 20 aaRS-depleted PURE systems by supplying 20 aaRSs from DNA through a 2.5-fold serial dilution for 20 cycles.

## Results

### Assay of translation efficiency and activities of 20 aaRSs

To systematically achieve regeneration of 20 aaRSs, we first evaluated the translation efficiency and specific activity of the 20 aaRSs expressed in the PURE system. In contrast to translation, which has been measured previously (*43*), the specific activity of each aaRS per protein has not yet been measured. The assay procedure is outlined in Fig. 1A.

**Fig. 1.**
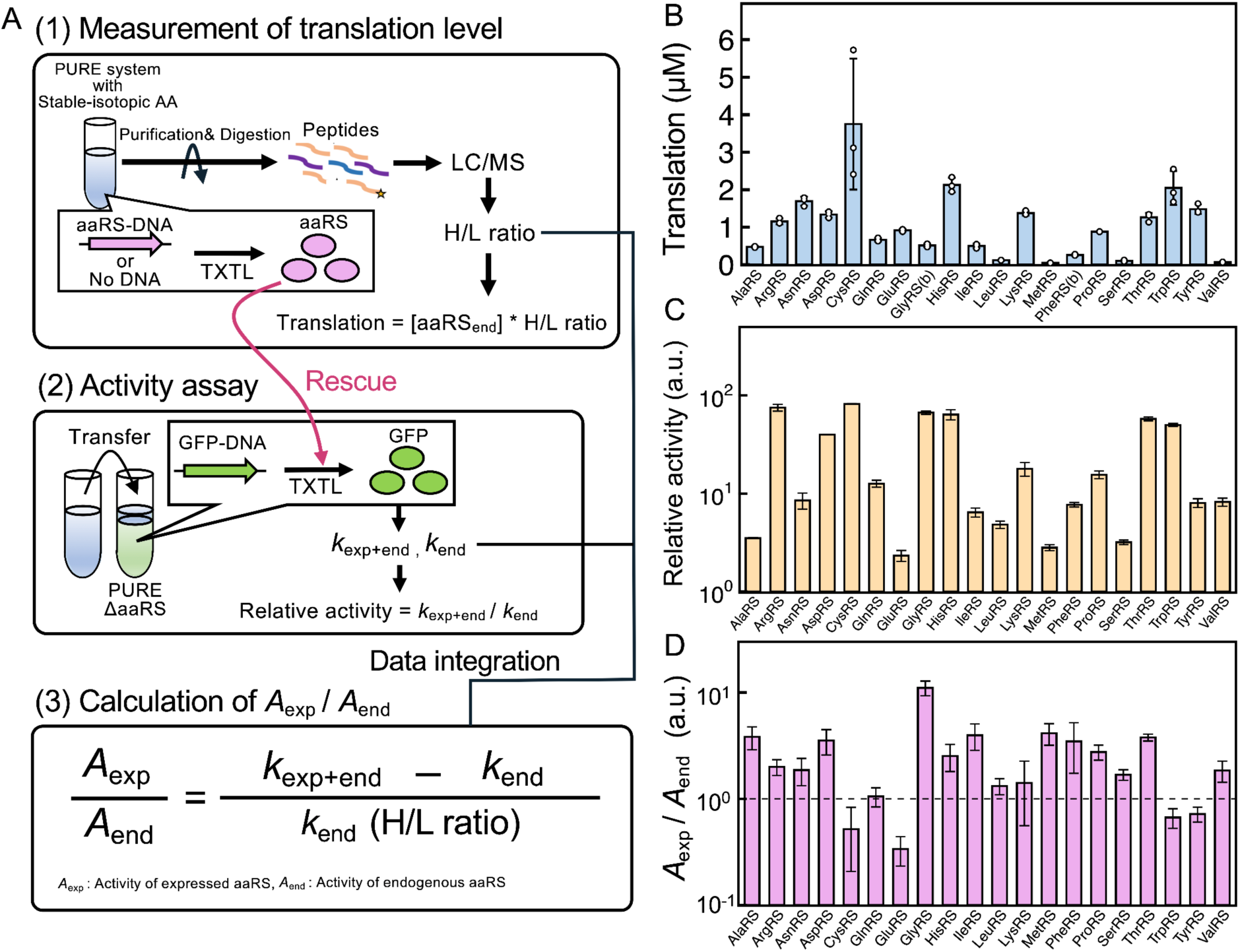
Characterization of 20 aaRS expressed in the PURE system. (A) Characterization scheme for 20 aaRS. Each aaRS was expressed in a customized 1^st^ PURE system (version 1) containing stable isotope-labeled amino acids at 30°C for 8 h. The expressed aaRSs were purified together with endogenous aaRSs, treated with trypsin, and analyzed using LC-MS to determine the heavy/light (H/L) ratio. The translation efficiencies were obtained by multiplying the H/L ratio by the known aaRS concentrations in the PURE system. AaRS activity was measured by expressing GFP in the 2^nd^ PURE system lacking the aaRS of interest and by assessing the GFP synthesis rate. The specific activity ratio (*A*_exp_/*A*_end_) was calculated from the H/L ratio and aaRS activity according to the equation. (B) Translation efficiency of each aaRS. Each point represents an individual peptide. Error bars represent the standard deviation. The H/L ratios are presented in Fig. S1. (C) The total activity of each aaRS. *k*_end_ represents the total activity of each endogenous aaRS and *k*_exp+end_ represents the total activity of both endogenous and expressed aaRSs. Error bars represent standard errors. The raw data for each aaRS are presented in Fig. S2-3. (D) Specific activity (aaRS activity per protein) ratio of expressed to endogenous aaRSs. Error bars indicate standard error.

For the translation efficiency assay (Fig. 1A (1)), the DNA encoding each aaRS (aaRS-DNA) was incubated with a customized PURE system (version 1) containing stable isotope-labeled amino acids (Lys and Arg) to label the newly synthesized aaRS. All the proteins, including the expressed aaRSs, were purified and digested with trypsin. The resulting peptides were quantified using liquid chromatography-mass spectrometry (LC-MS). The expressed aaRS, containing isotope-labeled amino acids, had a higher molecular weight (i.e., heavy) than the endogenous (i.e., light) aaRS in the PURE system, allowing for calculation of the abundance ratio (heavy to light (H/L) ratio) between the expressed and endogenous aaRSs (Fig. S1). Translation efficiency was determined by multiplying the H/L ratio by the known concentration of endogenous aaRSs (Fig. 1B). The translation efficiency varies greatly from 0.05 µM to 3.75 µM with the average of approximately 1 µM. Some aaRSs (LeuRS, MetRS, PheRS, SerRS, and ValRS) exhibited significantly lower translations than average, suggesting that there is room for improvement in these aaRS translations.

For the assay of the specific activity of the expressed aaRSs, each aaRS-DNA or no DNA (as a control) was incubated in the customized PURE system (version 1) using the same method as the translation assay. The mixture was then diluted with the customized PURE system lacking the aaRS of interest at various ratios, and GFP expression was monitored (Fig. 1A (2)). The increase in GFP fluorescence was plotted against dilution rate (Fig. S2) and the slopes of the regression curves were obtained. The slope of the data with aaRS-DNA represents the sum of the expressed and endogenous aaRS activity (*k*_exp+end_). The slope of the data with no DNA represents endogenous aaRS activity (*k*_end_). From these data, we obtained the relative activities (*k*_exp+end_/*k*_end_) of the total expressed aaRS to the total endogenous aaRSs (Fig. 1C), which also exhibited varying amounts. For TrpRS, we employed phi29 DNA polymerase instead of GFP as a reporter gene because GFP has only one tryptophan residue (Fig. S3) (*43*).

To estimate the specific activity of each expressed aaRS, the relative activity ((*k*_exp+end_ − *k*_end_) / *k*_end_) was divided by the relative translation amount (H/L ratio) obtained previously (Fig. 1A (3)). The resultant value (*A*_exp_/*A*_end_) represents the relative specific activity (per protein) of the aaRS expressed in the PURE system compared to that of the endogenous purified aaRS protein. The relative specific activities of most aaRSs were more than one (Fig. 1D), indicating that the expressed proteins of most aaRSs had similar or higher specific activities than the endogenous ones. This result suggests that sustainable regeneration can be achieved with lower protein expression for these aaRSs. However, some aaRSs (CysRS, GluRS, TrpRS, and TyrRS) exhibit lower specific activity than endogenous aaRSs, indicating room for improvement.

### Improvement of the 5’-terminal sequences

First, we sought to enhance the translation efficiency. The aaRS sequences used in this study originated from plasmids used for expression in *E. coli* (*44*). By comparing the 5’-untranslated region (UTR), we found that some aaRSs (LeuRS, MetRS, SerRS, and ValRS), which show relatively smaller translation in Fig. 1B, have a shorter 5’-UTR that lacks a spacer sequence and an epsilon sequence (Table S1), which are important regions for translation efficiency (*45*). We also found that for some aaRSs (LysRS, ProRS, GlnRS, LeuRS, and SerRS), the GC ratio up to the seventh residue, which should be as low as possible for translation (*46*), could be reduced without changing the amino acid sequences (Table S1). Accordingly, we added the 5’-UTR sequences, reduced the GC ratio in the N-terminal regions of aaRSs, and observed some improvements in translation efficiency (Fig. 2A).

**Fig. 2.**
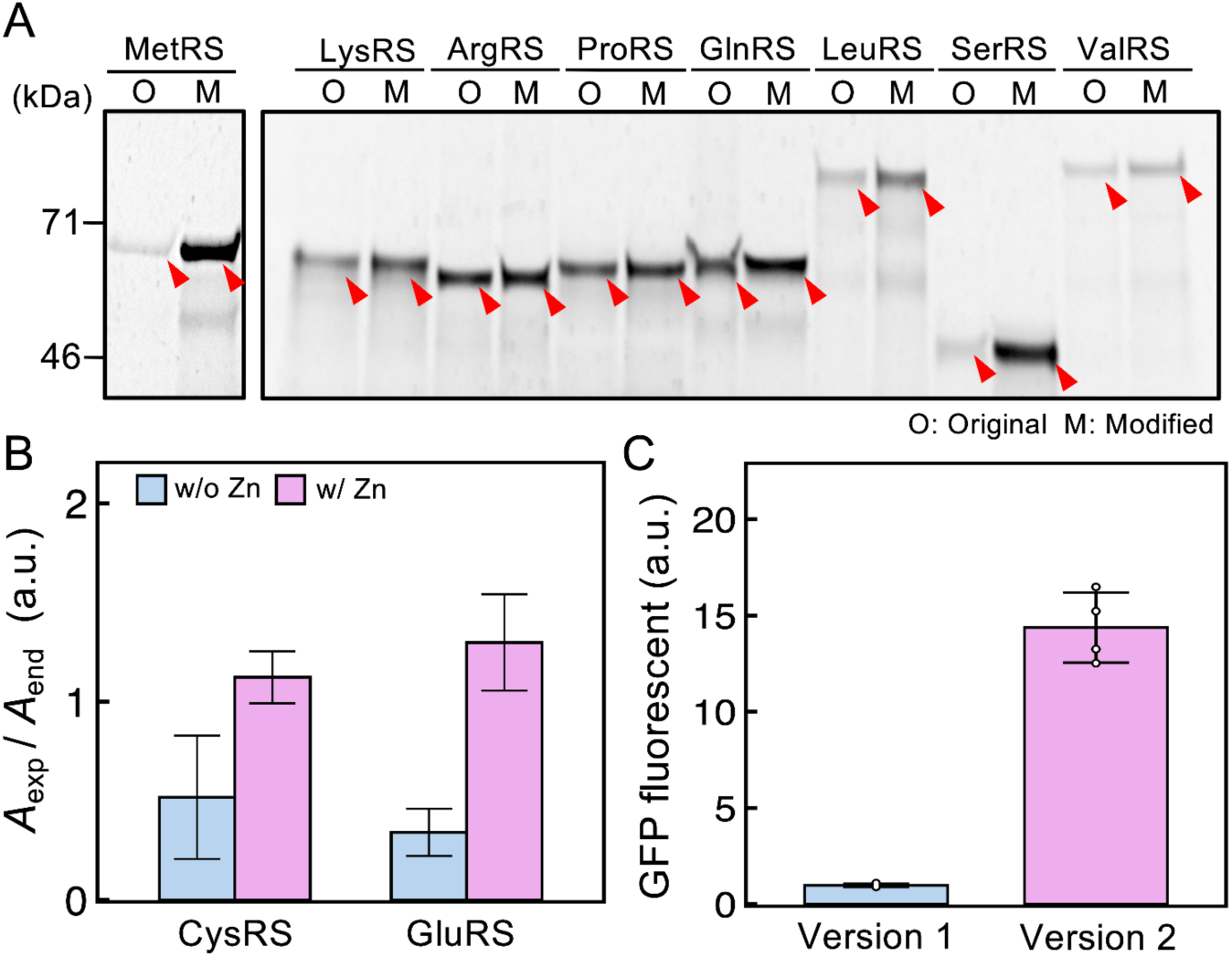
Improvement of 5’-terminal sequences and composition of PURE system. (A) Effect of the 5’-terminal sequence modification of the eight aaRSs, which exhibited lower translation efficiency. Each aaRS was expressed from each aaRS-DNA (5 nM) at 30 °C for 8 h in a customized PURE system (version 1) containing fluorescently labeled lysyl-tRNA. The mixture was subjected to SDS-PAGE and fluorescence was detected. The expected bands of each aaRS are indicated by the red arrowheads. The results for the original (O) and modified (M) sequences are presented. (B) Improvement in aaRS activity with zinc addition. Assays were performed in the same manner as in Fig. 1, with and without 1 mM Zn(OAc)_2_, and aaRS activity was calculated. Error bars represent the standard errors. (C) Improvement of the PURE system composition. The concentrations of non-protein components and T7 RNA polymerase in the PURE system were changed (Table S2), and translation activity was measured by GFP expression. Error bars represent the standard deviation of four independent experiments.

### Improvement of the composition of the PURE system

Two aaRSs (CysRS and GluRS), which exhibit low specific activities, contain Zn ions in their active sites (*47*, *48*). To address the possibility of zinc deficiency in these aaRSs, 1 mM of Zn(OAc)_2_ was added to the PURE system (version 1). The specific activities (*A*_exp_/*A*_end_) of CysRS and GluRS increased 2-4 times and reached one (Fig. 2B), indicating that the resultant specific activities were comparable to those of the endogenous aaRSs.

Subsequently, we changed the composition of the PURE system to improve its translation capacity. In a previous study (*43*) and in the experiments described so far, we used a customized PURE system tailored for DNA replication (termed “version 1” in this study), in which some components, such as T7 RNA polymerase, NTPs, and tRNA, were reduced (Table S2, version 1). By restoring the concentrations of these components to their original levels (*49*), the translation level increased by approximately 14-fold (Fig. 2C and Table S2). The PURE system of this composition was termed “version 2” in this study.

### Highly active aaRSs derived from other bacteria

One possible strategy for aaRS regeneration is to minimize the required translation level by enhancing the specific activities of the expressed aaRSs. We first focused on AlaRS, which is the most abundant aaRS in PURE systems. We selected AlaRSs from ten bacteria with growth rates similar to that of *E. coli* (*50*) (Fig. 3A). They were expressed in the PURE system and their activities were assessed using the GFP expression assay shown in Fig 1A(2)). Although all ten variants exhibited lower translation efficiencies than *E. coli* AlaRS (Fig. 3B), seven of which showed higher total activities (Fig. 3C). Among these, we selected the most active AlaRS (#4) from *G. stearothermophilus* (GsAlaRS) for subsequent regeneration experiments. Next, we investigated activities of other aaRSs isolated from *G. stearothermophilus*. Of the nine tested aaRSs, SerRS (GsSerRS) was approximately six times more active than *E. coli* SerRS (EcSerRS) when expressed in the PURE system (Fig. 3D). We also used GsSerRS for the subsequent regeneration experiments.

**Fig. 3.**
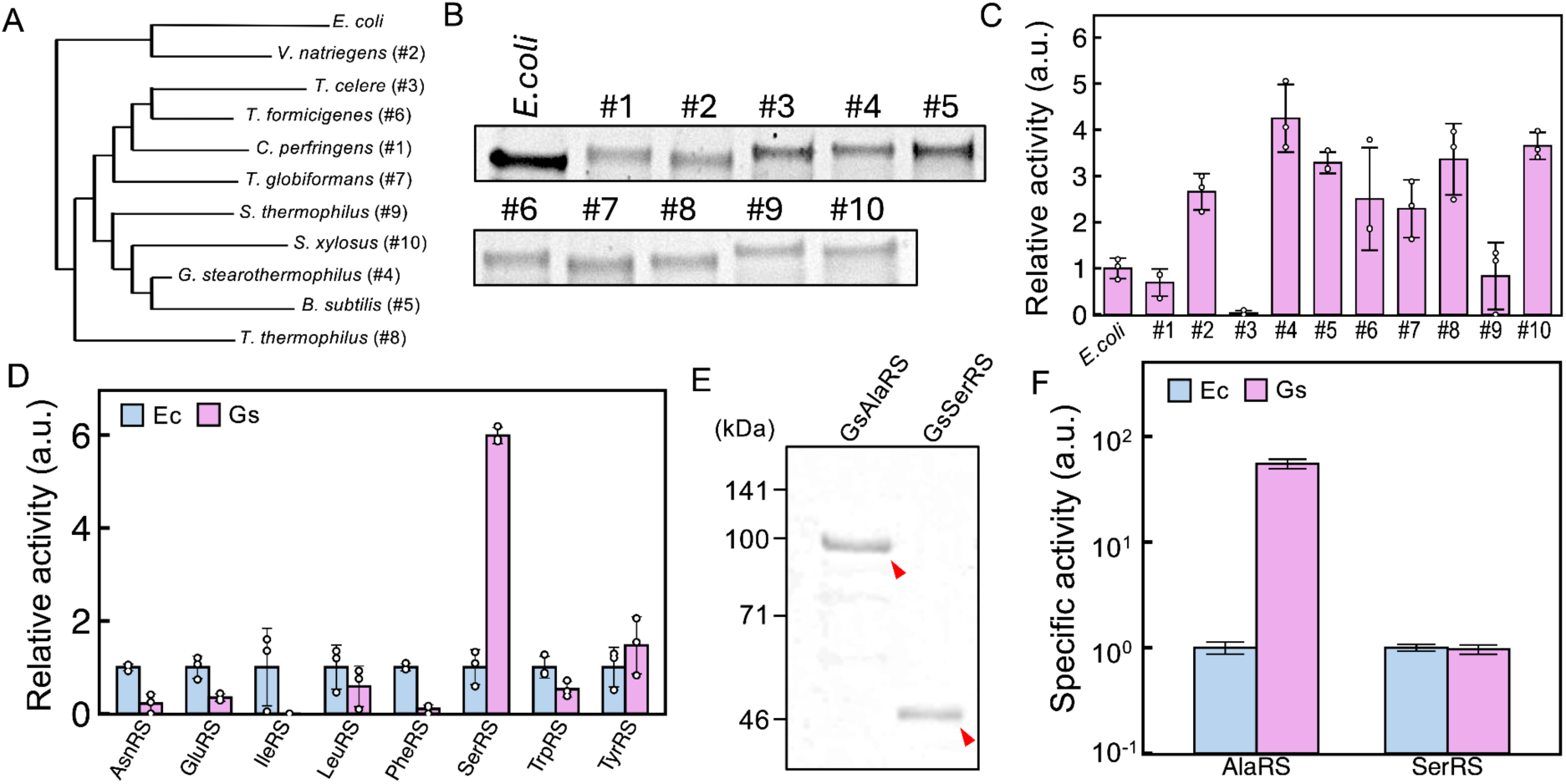
Translation and activities of aaRS derived from other bacteria. (A) Phylogenetic tree of AlaRSs obtained from *E. coli* and 10 other bacteria. A tree was constructed using the neighbor-joining method. (B) Translation of the AlaRSs. Each aaRS was expressed in the PURE system (version 1) with fluorescently labeled lysyl-tRNA at 30°C for 8 h. The mixture was analyzed by SDS-PAGE and fluorescence was detected. The entire gel image is shown in Fig. S4A. (C) Activity of AlaRSs. AlaRSs were expressed in the PURE system (version 1), diluted with the PURE system lacking AlaRS, and GFP fluorescence was measured as the AlaRS activity. Values relative to *E. coli* AlaRS are shown. Error bars show the standard deviations from three experiments. (D) Activity of eight aaRSs of *G. stearothermophilus*. Eight Gs-derived aaRSs were expressed and measured as described in (C). Error bars show standard deviations from three experiments. The results of SDS-PAGE analysis are shown in Fig. S4B. (E) SDS-PAGE analysis of recombinant GsAlaRS and GsSerRS purified from *E. coli*. (F) Specific activity of purified aaRS. Each purified aaRS was added to a PURE system lacking the target aaRS, and GFP expression was used to calculate specific activity. Error bars indicate standard error. The raw data are shown in Fig. S5.

To compare the specific activities of GsAlaRS and GsSerRS to *E. coli* ones, we purified recombinant GsAlaRS and GsSerRS from *E. coli* (Fig. 3E) and assayed for their activities. We found that GsAlaRS was approximately 50 times more active than its *E. coli* counterpart, whereas GsSerRS showed no significant difference (Fig. 3F). These results indicate that GsAlaRS is more active than EcAlaRS even when expressed in *E. coli*, whereas GsSerRS is more active only when expressed in the PURE system.

### Serial-dilution experiments for 1-10 aaRS regeneration

We attempted to regenerate 1-10 aaRSs by serial dilution experiments using the data and the improved aaRSs obtained above. The experimental procedure is illustrated in Fig. 4A. In the first reaction (round 1), 1-10 aaRS-DNAs of interest and firefly luciferase-encoding DNA (Fluc-DNA) were incubated in the customized PURE system (version 2), containing all 20 aaRS proteins, for 8 h at 30°C. The concentration of each aaRS-DNA was determined based on the translation efficiency and the specific activity ratio (*A*_exp_/*A*_end_) to regenerate the original aaRS ratios in the PURE system (Table S3). During incubation, aaRSs of interest and reporter luciferase were expressed. Thereafter, an aliquot of the reaction mixture was diluted 5-fold with another customized PURE system (PUREΔaaRS) that lacks each aaRS of interest, contains aaRS-DNA and Fluc-DNA at the same concentration, and incubated again. In this second reaction, both luciferase and the target aaRSs were continuously expressed if the lacking aaRS protein was sufficiently supplied (i.e., rescued) from the previous round of reaction. This serial dilution process was repeated for up to ten rounds, and at the end of each round, luciferase activity was assayed as a measure of translation activity. If each aaRS is regenerated continuously in sufficient amounts, translation activity should remain constant. As a negative control, we conducted the same experiment using DNA encoding GFP (GFP-DNA) instead of aaRS-DNA. Additionally, as a positive control, we conducted the same serial dilution experiment using GFP-DNA and a customized PURE system containing all 20 aaRS proteins (Full PURE).

**Fig. 4.**
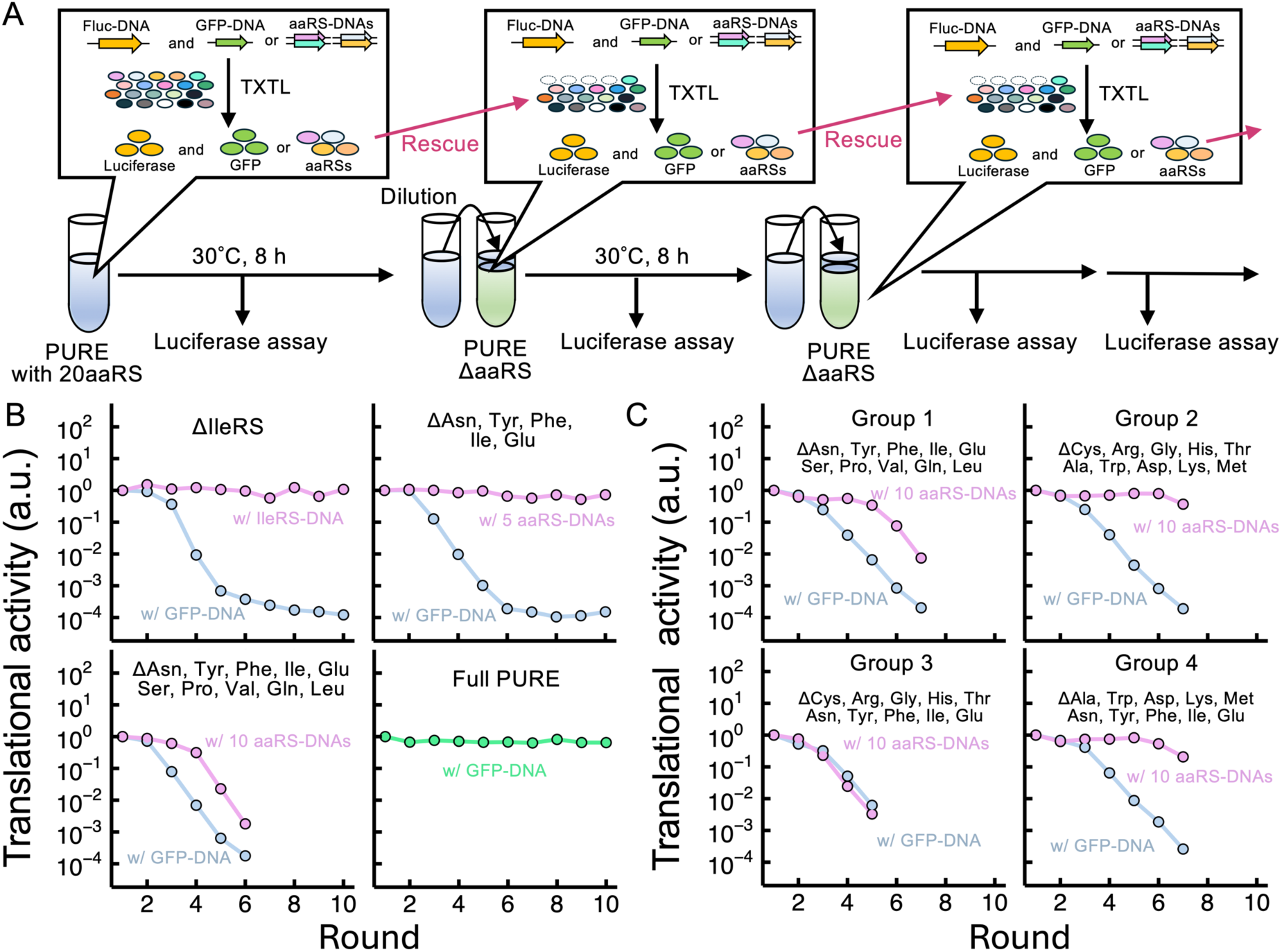
Serial dilution experiments for 1‒10 aaRS regenerations. (A) Scheme of serial dilution experiment. The first reaction mixture contains 1-10 aaRS-DNAs (5 nM in total) and 0.01 nM Fluc-DNA encoding luciferase in the customized PURE system (version 2) containing all 20 aaRS proteins. In the control experiment, GFP-DNA (5 nM) was used instead of 20 aaRS-DNA. Each DNA concentration is presented in Table S3. In the first reaction (round 1), aaRS, GFP and Luciferase were expressed at 30 °C for 8 h. From round 2 onwards, the reaction solution of the previous round was diluted 5-fold with the customized PURE system (version 2), which lacks each aaRS of interest (PUREΔaaRS) and contains the same DNA sets as round 1. The diluted mixture was then incubated at 30 °C for 8 h. If aaRSs are sufficiently synthesized in the previous round, the lack of aaRSs in the next round is rescued, and luciferase and aaRS are expressed again. Luciferase activity (luminescence) was measured after each round to assess the translation activity. (B-C) Trajectories of the translation activity in the serial dilution experiment. The translational activity in each round was plotted after normalization based on the value at round 1.

We observed that for a single aaRS (IleRS), five aaRSs (AsnRS, TyrRS, PheRS, IleRS, and GluRS), and the positive control (Full PURE), the translational activities at each round were maintained at a level similar to the initial level, at least until round 10 (Fig. 4B, upper and lower right panels). In contrast, the translational activity of the 10 aaRS-DNAs decreased continuously in each round (Fig. 4B, lower left panel). We conducted the same experiments using 10 aaRS-DNAs in different combinations. All translation activities eventually decreased (Fig. 4C). These results demonstrate that the current reaction conditions are insufficient for the regeneration of 10 aaRSs. This result is unexpected because according to the calculation based on the data obtained above, a sufficient amount of all aaRSs would be expressed. This implies that unknown factors diminish the expressions or activities.

### Inhibitory effect of some aaRS for translation

One of the possible factors that has not been considered in the previous experiment is the interaction among the 20 aaRS DNAs, because we measured each aaRS translation independently. To examine the effect of each aaRS DNA on the expression of other genes, we incubated each aaRS-DNA with Fluc-DNA in the PURE system and compared the luminescence produced by the expressed luciferase. The results were then plotted against the expected translation of each aaRS used in Fig. 1B. If there is simple competition between aaRS-DNA and Fluc-DNA for translation resources, luminescence should linearly decrease as the expected translation increases. As expected, approximately half of the aaRSs followed this trend (Fig. 5A, indicated by the wide grey line). In contrast, other eight aaRS-DNAs (GsAlaRS, AspRS, GlnRS, GluRS, GlyRS, IleRS, LeuRS and ValRS) are accumulated in the lower left part the figure (yellow), implying that these aaRS-DNA inhibit luciferase expression more than expected from simple competition.

**Fig. 5.**
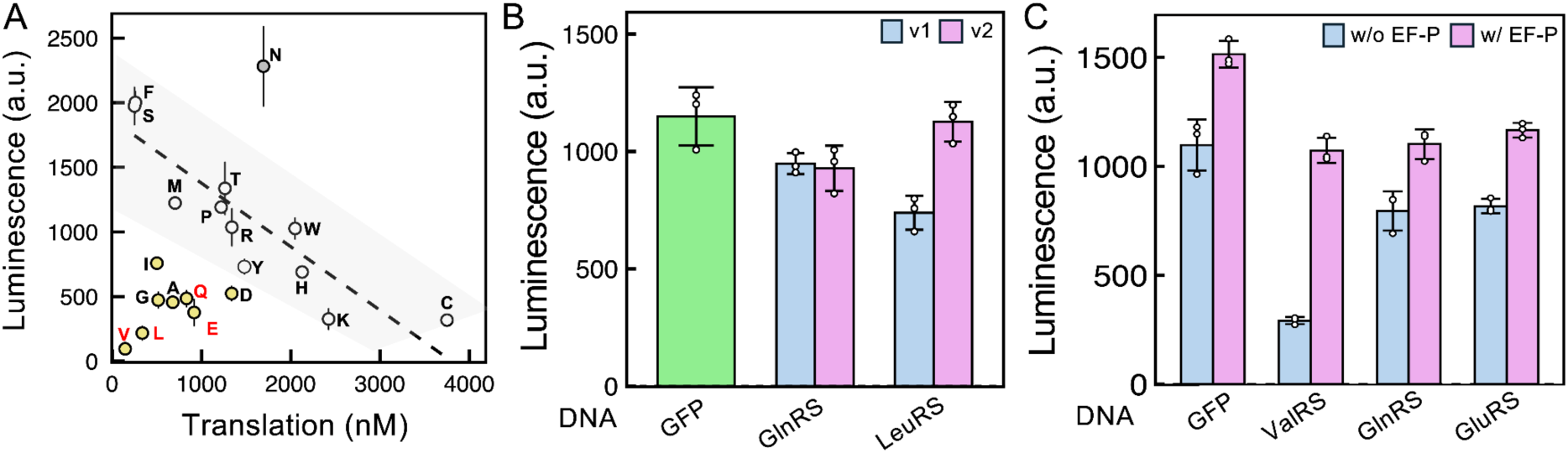
Inhibitory effect of each aaRS-DNA on translation of another gene. (A) Relationship between the translation of each aaRS (Fig. 1B) and co-expressed luciferase (luminescence). Each aaRS-DNA (5 nM) and Fluc-DNA (0.01 nM) were incubated in the PURE system (version 2) at 30°C for 8 h, and luminescence was measured. Each point represents an aaRS. The points that follow a linearly decreasing trend determined by the random sample consensus method are shown in white. Yellow points represent candidates for inhibitory aaRSs. (B) Effect of rare codons in GlnRS-DNA or LeuRS-DNA on inhibitory effect. One and four rare codons in GlnRS and LeuRS were substituted with the common codons in GlnRS_v2 and LeuRS_v2, respectively. Each aaRS-DNA (1 nM) was co-expressed with GFP-DNA (4 nM) and Fluc-DNA (0.01 nM) in the PURE system (version 2) at 30 °C for 8 h, and luminescence was measured. Error bars represent the standard deviation of three independent experiments. (C) Effect of EF-P on inhibitory effects. As in (B), each of the three aaRS-DNAs was co-expressed with GFP-DNA and Fluc-DNA, with or without the addition of EF-P (1 µM), and luminescence was measured. Error bars represent the standard deviations of three independent experiments.

Next, we attempted to eliminate the inhibitory effects of four aaRS-DNAs (LeuRS, ValRS, GlnRS, and GluRS) that require relatively large amounts for sustainable regeneration. One possible mechanism for translational inhibition is ribosome sequestration caused by ribosome stall. Inhibitory aaRSs may have a sequence that induces the stalling of ribosomes, which could reduce active free ribosomes and inhibit the translation of other genes. LeuRS contains four closely located rare codons (AGA, ATA, CGA, and CTA codons at positions 6, 9, 46, and 47, respectively) that may induce ribosome stalling at this position. Therefore, we created a modified LeuRS (LeuRS_v2) by replacing all rare codons with a common one, and compared its inhibitory effect when co-expressed with Fluc-DNA. Co-expression with the original LeuRS reduced luciferase activity by 40% compared to GFP-DNA, but this reduction was mitigated with LeuRS_v2 (Fig. 5B). GlnRS also contains a rare codon (AGA at position 6), while the substitution of this rare codon (GlnRS_v2) did not mitigate the inhibition effect (Fig. 5B). These results indicated the importance of avoiding multiple or clustered rare codons.

The three aaRSs (ValRS, GlnRS, and GluRS) do not contain a region with multiple rare codons, but commonly contain proline repeats. GlnRS and GluRS have a proline doublet (PP), while ValRS has a proline triplet (PPP), which is known to cause ribosome stalling (*51–53*). Ribosome stalling at the polyproline site can be resolved by addition of EF-P (*52*, *53*). We tested whether EF-P could eliminate the inhibition caused by these three aaRSs. When GlnRS, GluRS, and control GFP (no polyproline) were co-expressed with Fluc-DNA, EF-P addition slightly increased luciferase luminescence, even for GFP, whereas when ValRS was co-expressed, EF-P significantly enhanced luminescence (Fig. 5C). This suggests that the inhibitory effect of ValRS, which could be caused by its proline triplet, can be alleviated by the addition of EF-P.

In these experiments (Fig. 5B and 5C), we identified two methods (using LeuRS_v2 and adding EF-P) to mitigate the inhibitory effects of LeuRS and ValRS-DNAs. To verify the total effect of these methods under conditions that can be used for 20 aaRS regeneration experiments, we performed a co-expression experiment of Fluc-DNA with all four inhibitory aaRS-DNAs at realistic concentrations. We observed approximately 40% inhibition of luciferase activity, which was reduced to the original level by LeuRS_v2 and the addition of EF-P (Fig. S6).

Another interesting result shown in Fig. 5A is the higher luminescence when co-expressed with AsnRS-DNA (N) than the expected competition line. This result suggests that AsnRS expression increases the translation of other genes. This result implies that AsnRS protein is insufficient in the PURE system. This point was not addressed in this study, but should be improved in the future.

### Calibration of DNA concentration ratio

Another possible factor that could cause the failure of aaRS regeneration is the deviation from the expected expression level when multiple aaRSs are co-expressed owing to the competition of transcription/translation resources among aaRS-DNAs. In the previous experiment in Fgi. 4, we prepared an aaRS-DNA mixture that was expected to express 20 aaRS at the optimum balance based on the individually measured data (Table S4, left). In this study, we prepared a 20 aaRS-DNA mixture using the same method and measured its expression in a PURE system (version 2), as shown in Fig. 1A(1)), using isotope labeling and LC-MS. The actual translation of each aaRS (Fig. 6A), and the ratio of the actual to predicted translation (Fig. 6B) were calculated. If each aaRS is expressed as predicted, the ratio should be one for all aaRSs. However, Fig. 6B shows that the ratio (TL/Predict) varies from 0.04 to 1.2 (standard deviation is 0.34), and also the average ratio (0.37) was smaller than one, indicating that when co-expressed, both the balance and the averaged translation level of 20 aaRS are different from our prediction.

**Fig. 6.**
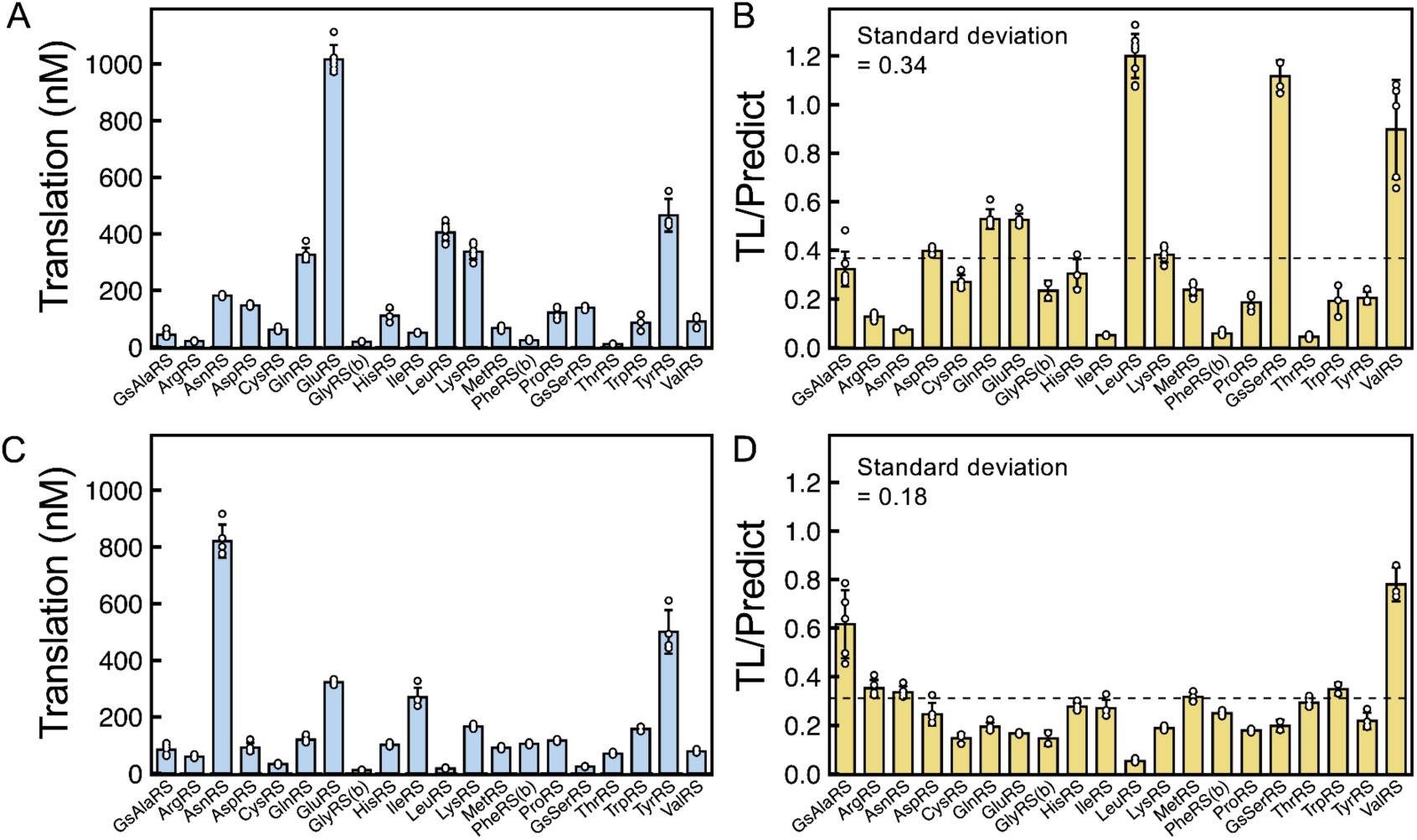
Quantification and calibration of the co-expressed 20 aaRSs. All 20 aaRSs were incubated in a customized PURE system (version 2) containing stable isotope amino acids at 30 °C for 8 h. After LC-MS analysis using the method shown in Fig. 1A(2), the translation of each aaRS-DNA was calculated. (A, B) Results with the aaRS-DNA composition (Table S4, left, 5 nM in total), which was determined based on the individually measured translation efficiency in Fig. 1B. The translation amount of each aaRS is shown in (A), and the ratio of the translation to the prediction from the individually measured data is shown in (B). (C, D) Results with the aaRS-DNA composition (Table S4, right, 2 nM in total), which was calibrated based on the deviation from the prediction shown in (B). The translation amount of each aaRS is shown in (C), and the ratio of the translation to the prediction after the calibration is shown in (D). Each point represents an individual detected peptide, with error bars indicating standard deviation. H/L ratio plots are shown in Fig. S7. The dotted line represents the average TL prediction.

To calibrate the expression balance, we adjusted the DNA concentration ratios by increasing the DNA for insufficient aaRSs and decreasing it for sufficient aaRSs based on the data in Fig. 6B (Table S4, right). Additionally, to increase the total expression level, we used LeuRS_v2-DNA, which omitted rare codons, and added EF-P to mitigate the inhibitory effects of LeuRS and ValRS-DNA. After this calibration, the translation level (Fig. 6C) and the ratio to the predicted level (Fig. 6D) were measured by the same method. The standard deviation of the ratio (0.18) became smaller than the value before calibration, indicating that the expression balance was improved. Although the average ratio (0.31) still did not approach one, we decided to conduct the regeneration experiment because the expected values were determined based on the aaRS concentrations of the original PURE system, which are usually excessive.

### Serial-dilution experiments for 20 aaRS regeneration

Finally, we attempted to regenerate 20 aaRSs using all the improvements we had shown above. The experimental procedure is shown in Fig. 7A, which is almost the same as Fig. 4A, but the reaction mixture contains all 20 aaRS-DNAs, and the PURE system used for dilution (composition is shown in Table S5) lacks all 20 aaRSs. First, we performed serial dilution experiments at four total DNA concentrations (2, 3, 5, and 7.5 nM) to determine the optimal total DNA concentration. Translation activity was maintained at a similar level up to 10 rounds at 2 and 3 nM, whereas it gradually declined at 5 and 7.5 nM (Fig. S8), indicating that higher DNA concentrations were unsuitable for aaRS regeneration. We further conducted serial dilution experiments under four conditions (2 or 3 nM aaRS-DNA mixture and 2 or 2.5-fold dilution) for up to 20 rounds and found that the translation activities (luciferase activity) were maintained at a certain level in the final rounds under all four conditions (Figs. 7B and S9). These data suggested that all 20 aaRSs were sustainably regenerated during the serial dilution cycle.

**Fig. 7.**
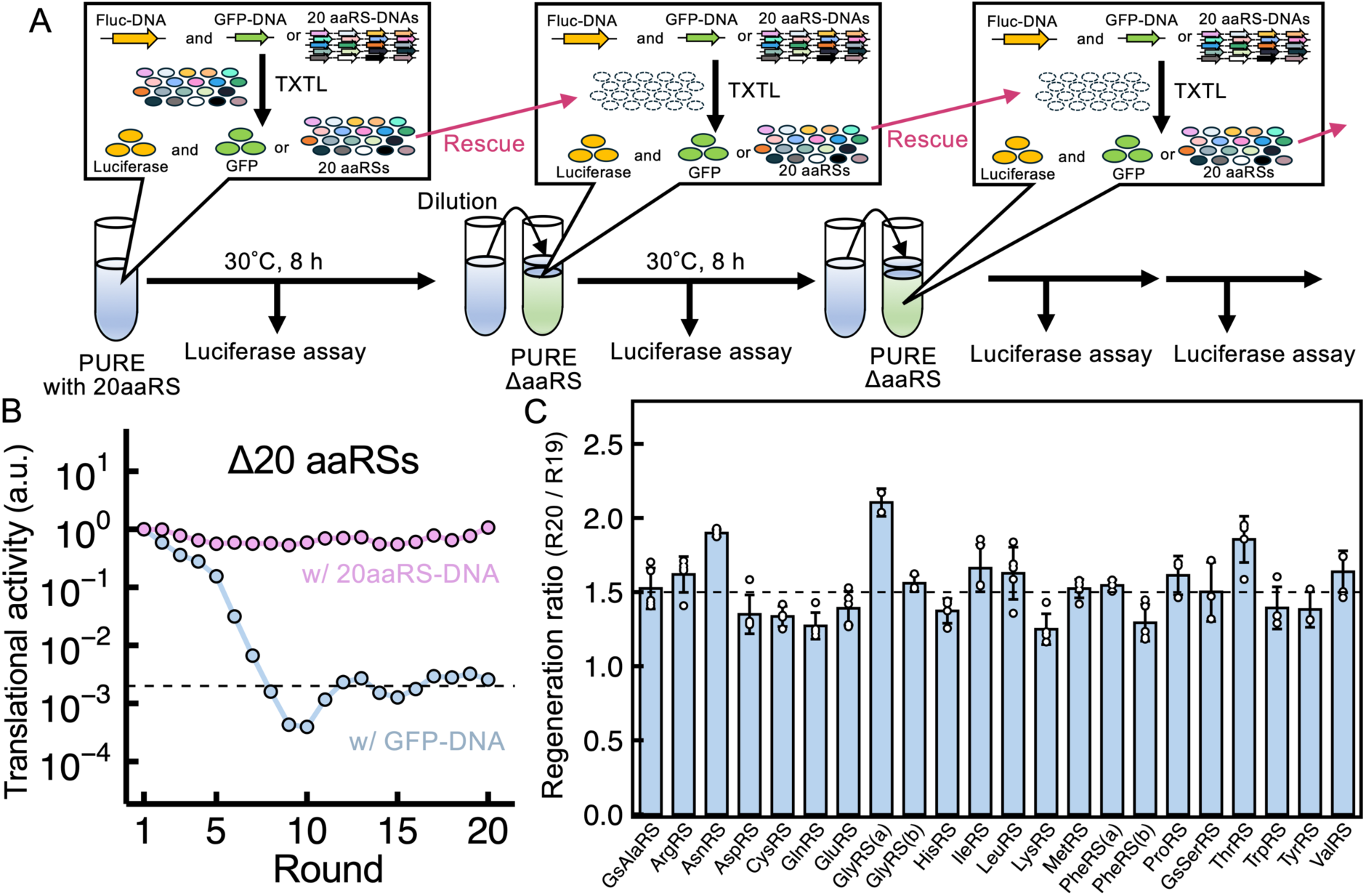
Serial Dilution experiments for 20 aaRS regeneration. (A) Scheme of the serial dilution experiment. The first reaction mixture contained 20 aaRS-DNAs (2 nM in total) and 0.01 nM Fluc-DNA encoding luciferase in the customized PURE system (version 2) containing all 20 aaRS proteins. In the control experiment, GFP DNA (2 nM) was used instead of the aaRS-DNA. At round 1, aaRS, GFP, and luciferase were expressed. From rounds 2 to 20, the reaction solution of the previous round was diluted 2.5-fold with a customized PURE system that lacked all 20 aaRS proteins (PUREΔaaRS) but contained 20 aaRS-DNA (or GFP-DNA) and Fluc-DNA at the same concentrations. Then, the reaction mixture was incubated. If aaRS is sufficiently synthesized in the previous round, the lack of each of the 20 aaRSs is rescued, and thus luciferase and aaRS are expressed again. Luciferase activity (luminescence) was measured after each round to assess translation activity. In round 20, stable isotope-labeled amino acids were added to the mixture to measure the aaRS translation. (B) Trajectories of the translation activity. The translation activity (luciferase activity) was normalized based on luminescence after round 1. (C) Regeneration ratio of each aaRS at round 20. The ratio of isotope-labeled aaRS (heavy) to unlabeled aaRS (light) was measured using LC-MS. The heavy and light peptide concentrations corresponded to aaRSs synthesized at round 20 and those synthesized until R19, respectively. If each aaRS concentration is maintained, the ratio should be near 1.5 at this dilution rate. Each point represents an individual detected peptide, with error bars showing standard deviation.

To further verify the sustainable regeneration of 20 aaRSs, we compared aaRSs expressed at round 20 to those expressed before round 20. We first replaced the amino acids in PUREΔaaRS with stable isotopes at round 20 and quantified the synthesized 20 aaRSs using LC-MS. We then calculated “the regeneration ratio of each aaRS (R20/R19)”, which represents the ratio of aaRSs expressed at round 20 to aaRSs transferred from round 19, from the H/L ratio by normalizing the isotope ratio. If 20 aaRS was expressed at round 20 to the same level as in round 19 (i.e., sustainably regenerated), the regeneration ratio should be 1.5, because the reaction mixture was diluted 2.5-fold in each round (1 for the transferred aaRS and 1.5 for the newly synthesized aaRS). The regeneration ratios of all 20 aaRSs were approximately 1.5 (Fig. 7C), indicating that all 20 aaRS were sustainably regenerated even at round 20, where the purified aaRS proteins included in the first reaction in round 1 must be diluted to approximately 10^−8^.

In this study, we employed six improvements: 1) sequence modification, 2) zinc addition, 3) PURE system composition change, 4) GsAlaRS and GsSerRS utilization, 5) diminishing inhibitory effects, and 6) adjustment of DNA concentration for co-expression. To evaluate the contribution of each improvement to regeneration, we conducted serial dilution experiments again by restoring each improvement. We observed an immediate decline in translation activity when any one of the five improvements was restored, except for 2) zinc addition (Fig. 8). These results demonstrate that all five improvements, excluding zinc addition, were crucial for the sustainable regeneration of 20 aaRSs.

**Fig. 8.**
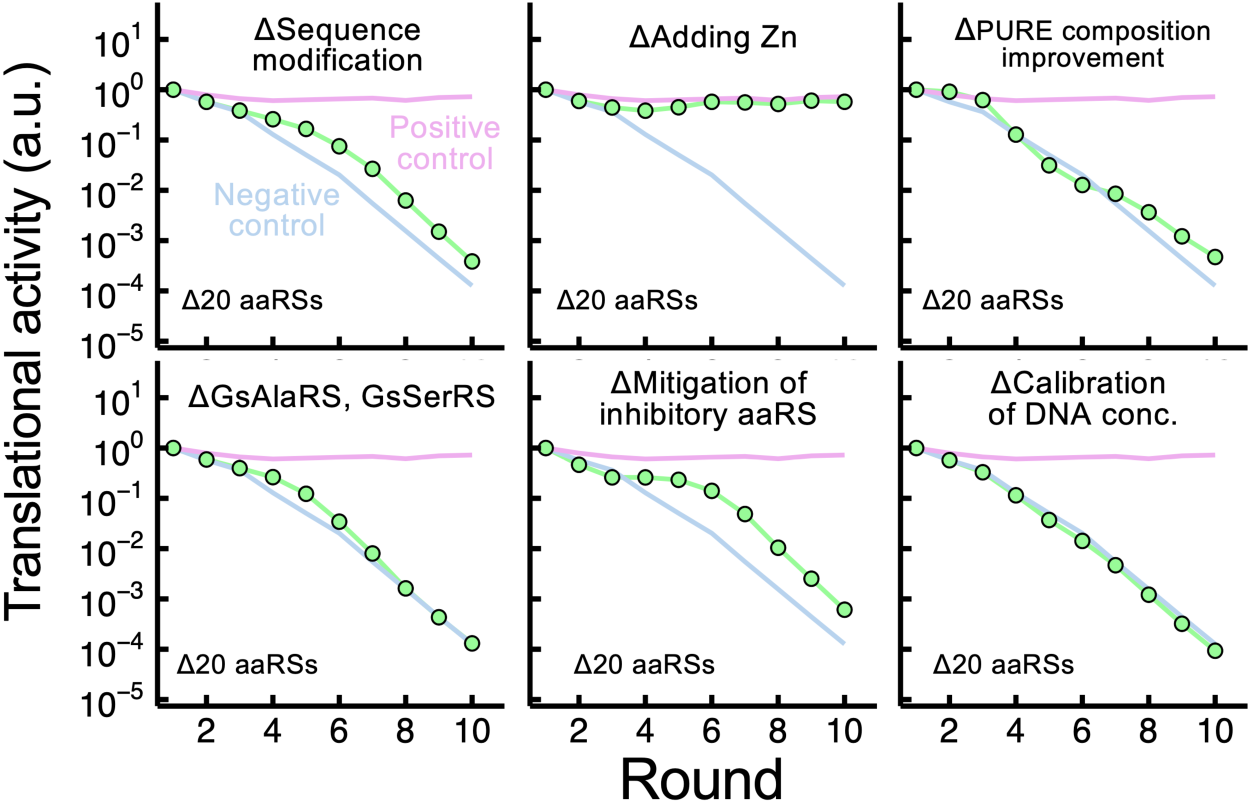
Contribution of each improvement to 20 aaRS regeneration. Serial dilution experiments, shown in Fig. 7A, were performed under conditions that reverted each of the six improvements. The trajectories of the translation activity are shown (green points). The red line indicates the condition in which all improvements were applied (positive control, the same condition as Fig. 7B), and the blue line indicates the condition in which GFP DNA was used instead of 20 aaRS-DNAs (negative control).

## Discussion

Self-reproduction is one of the most unique functions of living organisms, and its artificial realization is a significant challenge in bottom-up synthetic biology (*2*, *14*, *15*, *17*). To advance this challenge, we focused on 20 aaRSs, which comprise approximately half of the translation proteins included in the PURE system, except for ribosomes. Unlike previous studies, we measured the specific activities of all aaRSs expressed in the PURE system, together with translation, and systematically improved both translation efficiencies and specific activities of aaRSs with insufficient activities using six methodologies: 1) sequence modification, 2) zinc addition, 3) PURE system composition change, 4) GsAlaRS and GsSerRS utilization, 5) diminishing inhibitory effects, and 6) calibration of the DNA concentration ratio. Because of these improvements, we succeeded in the sustainable regeneration of all 20 aaRSs for up to 20 rounds. This result represents a significant advancement toward the realization of self-synthesizing artificial systems. In addition, the methodologies and knowledge obtained in this study will be useful for further increasing the regeneratable proteins in the future.

One of the new strategies employed in this study is to adjust the expression balance for 20 aaRSs based on translation efficiency as well as the specific activity of each aaRS. Our previous approach relied solely on translation efficiency and endogenous aaRS concentration to determine DNA ratios, resulting in an insufficient regeneration performance (*43*). In this study, we introduced a more refined strategy by integrating specific activities into the calculations. Accordingly, we found that some aaRSs (e.g., GlyRS, IleRS, and MetRS) were more active when expressed in the PURE system than the purified proteins, which means that the required expression level of aaRSs for regeneration is lower than previously thought. Another important insight obtained is that the DNA concentrations adjusted based on the individually measured data provided discrepancies between the predicted and actual translation amounts (Fig. 6B), probably due to the transcription/translation competition between genes, and calibration with the co-expression data was required for balanced expression. Even in the final condition, some aaRS expression levels (e.g., LeuRS) were still lower than predicted (Fig. 6D). Understanding the reason for this discrepancy will allow for more precise control of gene expression in the PURE system.

A novel finding of this study was the highly active AlaRS and SerRS from *G. stearothermophilus* when expressed in the PURE system. Generally, more active translation factors require fewer translation resources to achieve the same level of activity and are thus useful for self-regeneration. Although the exact mechanism by which these two aaRSs exhibit high activity in the PURE system remains unclear, the approach of selecting highly active translation factors from other bacteria can be a useful strategy to improve translation factors.

Another novel finding is the translation inhibition caused by some aaRS DNAs, such as LeuRS, ValRS, GlnRS, and GluRS (Fig. 5A). Considering that the inhibitory effects were mitigated by removing rare codons or adding EF-P, a possible mechanism of the inhibitory effect is ribosome stalling, which decreases the availability of ribosomes. Several ribosome-stalling sequences, other than rare codons or polyproline, have been reported (*54*, *55*). These stalling sequences may have caused the remaining inhibitory effects of GlnRS and GluRS. This problem may be serious when the number of co-expressed genes is increased because the inhibitory effect is expected to accumulate. Further identification of inhibitory sequences is required to realize self-reproducible systems. In addition, knowledge of ribosome stalling sequences will contribute to a deeper understanding of translational mechanisms in natural living systems.

Although this study achieved stable regeneration of 20 aaRSs, some challenges remain. The first is the regeneration of additional genes, including other translational proteins, ribosomes, and tRNAs. The major obstacle is the translation capacity of the current PURE system, which can express up to 3.8 mg/ml of proteins under dialysis conditions (*49*), which is still lower than the protein concentration in our customized PURE system (7.1 mg/ml, PURE ver 7). One possible solution to this problem is to use a highly diluted PURE system that contains fewer proteins, as proposed by Ganesh *et al.* (*56*). However, we think that the regeneration of more genes may be easier than we thought because the specific activities of many newly expressed aaRSs in the PURE system were higher than those of purified aaRSs (Fig. 1D), which means that we can maintain the same level of translation activity with fewer proteins when expressed endogenously than predicted. Another challenge for self-reproduction is coupling with DNA replication. As shown previously, the buffer composition optimized for translation is significantly different from that optimized for DNA replication (*57*). Further studies are needed to develop a system that combines high translational activity with DNA replication.

## Materials and Methods

### DNA preparation

The DNA encoding each aaRS was prepared as follows. First, we PCR-amplified the DNA fragments encoding each aaRS using primers 1 (GCGAAATTAATACGACTCACTATAGGG) and 2 (CCGCTGAGCAATAACTAGCATAACC) and the plasmid (pET-aaRS) as templates, prepared in our previous study (*43*). The PCR product was purified using the QIAquick PCR Purification Kit (QIAGEN), which was used for all DNA purification procedures in this study. Nine sequence-modified pET-aaRSs were prepared by PCR amplification followed by self-ligation using a mutated and In-Fusion cloning Kit (TaKaRa). The circular DNA encoding the phi29 DNA polymerase used in the aaRS activity assay (Fig. S3) has been described previously (*20*, *43*). The plasmid that encodes GFP under the T7 promoter (pET-GFP) was previously constructed as pETG5tag (*58*). A linear DNA fragment encoding GFP-DNA was prepared by PCR using pET-GFP as a template and primers 1and 2. A plasmid encoding firefly luciferase under the T7 promoter (Fluc) was previously constructed (*59*). A linear DNA fragment encoding firefly luciferase (Fluc-DNA) was prepared by PCR using pET-Fluc as a template and the primers 3 (TTCTCTGCCCAATACGCAAAC) and 4 (GAGCAGACAAGCCCGTCAG). Linear DNAs encoding aaRS under the T7 promoter derived from other bacteria were prepared as follows. First, the amino acid sequences of the aaRS genes of the 10 bacteria were obtained from the NCBI database. After optimizing the nucleotide sequence for *E. coli* translation using three company sites or algorithms (Eurofin (Fig. 3B-C), and Twist Bio Sciences (Fig. 3D), and CodHonEditor (*60*) (Fig. 4-8), the gene sequence and upstream T7 promoter were synthesized and inserted into the pET vector using Twist Bio Science’s artificial gene synthesis service. The linear DNA fragment encoding each was prepared by PCR using each pET-aaRS as a template and primers 1 and 2. All plasmid sequences used in this study are shown in the Supplemental Information.

### PURE system preparation

The customized PURE system used in this study was prepared from independently purified components in our laboratory. Protein purification was conducted using affinity chromatography with a histidine tag, followed by gel-filtration chromatography as described previously (*61*). The compositions of the versions 1 and 2 are listed in Table S2. The final compositions used in Fig. 7 are listed in Table S5.

### Stable isotope-labeling and quantifying of PURE expressed aaRS

To measure the translation efficiency, aaRS-DNA (5 nM) was incubated at 30°C for 8 h in a customized PURE system in which Lys and Arg were replaced with stable isotope amino acids (0.36 mM ^13^C_6_ L-lysine-2HCl, 0.36 mM ^13^C_6_, ^15^N_4_ L-arginine-HCl) (Thermo Scientific) to express aaRS. Then, 10 µL of the reaction mixture was used to purify the proteins contained in the PURE system using a His-tag spin column (New England Biolabs), according to the manufacturer’s instructions.

The eluate was concentrated by methanol-chloroform precipitation and digested according to a phase-transfer surfactant-aided protocol (*62*). The pellets were dissolved in 20 µL buffer (10 mM sodium deoxycholate, 10 mM sodium N-lauroylsarcosinate, and 50 mM NH_4_HCO_3_), reduced with 10 mM tris(2-carboxyethyl)phosphine at 37°C for 30 min, alkylated with 20 mM iodoacetamide at 37°C for 30 min, quenched with 20 mM L-cysteine, and digested with 100 ng Lys-C/Trypsin Protease Mix (Thermo Scientific) at 37°C overnight. After digestion, the detergents were precipitated by adding final 1% trifluoroacetic acid (TFA), followed by centrifugation at 15,000 × g for 5 min at 4°C. The supernatant was desalted using GL-Tip SDB (GL Sciences) according to the manufacturer’s instructions and dried under reduced pressure. For the aaRS sample at round 20 (Fig. 7C), we used GL-Tip SDB-SCX (GL Sciences) instead of GL-Tip SDB for further fractionation to improve detection efficiency (*63*). In brief, peptides were concentrated on SDB-SCX and then eluted stepwise in the following buffers: 0.1% TFA, 0.1% TFA, 80% acetonitrile, 0.5% TFA, 30% acetonitrile, 1% TFA, 30% acetonitrile, 3% TFA, 30% acetonitrile, 3% TFA, 30% acetonitrile, 100 mM ammonium acetate, 4% TFA, 30% acetonitrile, 500 mM ammonium acetate, and 30% acetonitrile, 500 mM ammonium acetate. The eluates were dried under reduced pressure.

Dried peptides were dissolved in 0.1% TFA. The peptides were concentrated and separated using a nano-LC system (UltiMate 3000, Thermo Scientific, Bremen, Germany) equipped with a trap column (C18, 0.075 × 20 mm, 3 µm, Acclaim PepMap 100, Thermo Scientific) and nanocapillary analytical column (C18, 0.075 × 150 mm, 3 µm, Nikkyo Technos, Tokyo, Japan) at a flow rate of 300 nl/min. Mobile phases A (0.1% formic acid) and B (acetonitrile and 0.1% formic acid) were combined in the following gradient: 5% B for 5 min, 5–40% B for 75 min, 40–90% B for 1 min, and 90% B for 4 min. MS analysis was performed using an Orbitrap mass spectrometer (Q Exactive, Thermo Scientific) equipped with a nanospray ion source (Nanospray Flex, Thermo Scientific) with the following parameters: spray voltage, 2.2 kV, positive mode; scan range, *m/z* 310–2,000, and 70,000 resolution. The ten most intense multiply charged ions (*z* = 2–4) were fragmented in the collision cell by high-energy collisional dissociation (HCD) with a normalized collision energy of 30%. Calculation of peak areas and H/L ratio was performed using Skyline software (MacCoss group at the University of Washington, WA, USA).

In the quantification of aaRS regenerated in round 20 in Fig. 7C, we assumed that 40% of the regenerated aaRS was not labeled with stable-isotope amino acids because of the influence of the normal amino acids carried over from round 19, and the aaRS regenerated at round 20 was calculated by dividing the H/L ratio by 0.6.

### Activity assay of the aaRSs synthesized in the PURE system

First, the DNA encoding each aaRS (5 nM) was incubated at 30 °C for 8 h using a customized PURE system (version 1), in which Lys and Arg were replaced with stable isotope amino acids (Thermo Scientific) to express aaRS. Thereafter, an aliquot of the reaction solution was diluted with dilution buffer (50 mM HEPES-KOH (pH7.6), 100 mM KCl, 10 mM MgCl_2_, 7 mM 2-mercaptoethanol, 5 mg/mL BSA, 10 mM DTT, and 30% glycerol) at different rates and added to the second customized PURE system (version 1), which lacks aaRS of interest and contains GFP-DNA (3 nM). For AsnRS, CysRS, GluRS, HisRS, and TyrRS, Fluc-DNA (5 nM) was included as a translation load in the 2nd PURE system. The mixture was then incubated at 30 °C for 12 h. GFP fluorescence was measured every 15 min for up to 4 h and every 30 min thereafter (Mx3005P; Agilent Technologies). The slope of the GFP fluorescence from 0 to 4 h was evaluated as the GFP synthesis rate. The volume ratio of 1st PURE to 2nd PURE system was then plotted against the GFP synthesis rate, and the specific activities *k*_exp+end_ and *k*_end_ were calculated using linear regression. For TrpRS, the DNA replication rate was measured instead of the GFP synthesis rate as previously described (*43*).

The ratio of the specific activity of expressed aaRS to that of endogenous aaRS (*A*_exp_/*A*_end_) was calculated using the two obtained specific activities (*k*_exp+end_ and *k*_end_ obtained above) according to the following equation (Equation 1):

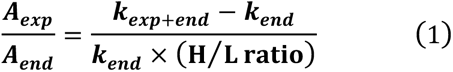

### SDS-polyacrylamide gel electrophoresis for quantifying aaRS expression amount

DNA (5 nM) encoding each aaRS was incubated at 30 °C for 8 h in a customized PURE system (version 1) containing fluorescently labeled lysyl-tRNA (FluoroTect GreenLys, Promega). After expression, an aliquot (5 µL) was treated with 0.5 µL of 5 mg/ml RNase A (QIAGEN, Hilden, Germany) at 37 °C for 30 min, incubated at 95 °C for 5 min in SDS sample buffer (17 mM Tris-HCl (pH 7.4), 0.7% SDS, 0.3 M 2-mercaptoethanol, and 3% glycerol) and subjected to 10% SDS-PAGE. The synthesized fluorescent-labeled proteins were detected using FUSION-SL4 (Vilber-Lourmat) and band intensities were analyzed using ImageJ.

### Activity assay of aaRS derived from other bacteria

As shown in Fig. 1, each aaRS was expressed in the 1^st^ PURE system, diluted with buffer, and added to the 2^nd^ PURE system lacking the aaRS of interest to monitor the GFP expression. GFP fluorescence was measured every 15 min up to 4 h and every 30 min thereafter (Mx3005P, Agilent Technologies). The slope of GFP fluorescence from 0 to 4 h was evaluated as the activity. To quantify the background activity of endogenous aaRSs, we used the PURE system without incubating any DNA, and the activity of each aaRS was evaluated by subtracting the background activity from the quantified activity of aaRS.

### Expression and purification of aaRS derived *G. stearothermophilus*

Expression plasmids encoding GsAlaRS (pET-GsAlaRS) and GsSerRS (pET-GsSerRS_v2) with histidine tags at the N-terminus and C-terminus, respectively, were synthesized by Twist Biotech (sequences are shown in Supplemental Information). The plasmids were introduced into Rosetta (DE3) pLysS (Novagen) and cultured at 37°C with shaking at 120 rpm in 1 L LB medium. In the late log phase, isopropyl *β*-d-thiogalactopyranoside (final concentration of 1 mM) was added and further incubated for 3 h. After harvesting, each aaRS was purified by affinity chromatography with a histidine tag, followed by gel-filtration chromatography using the same method as that used for the other protein factors of the PURE system, as described above (*61*). The purified Gs-derived aaRS band was subjected to SDS-PAGE and stained with Coomassie Brilliant Blue (Fig. 3E).

### Specific activity measurements of the purified GsaaRSs (Fig. 3F)

Purified AlaRS or SerRS derived from *E. coli* and *G. stearothermophilus* were diluted with dilution buffer (50 mM HEPES-KOH (pH7.6), 100 mM KCl, 10 mM MgCl_2_, 7 mM 2-mercaptoethanol, 5 mg/mL BSA, 10 mM DTT, and 30% glycerol) at some different rates and added to the customized PURE system that lacks AlaRS or SerRS and contains GFP-DNA (3 nM). The mixture was incubated at 30 °C for 12 h. GFP fluorescence was measured every 15 min for up to 4 h and every 30 min thereafter (Mx3005P; Agilent Technologies). The slope of the GFP fluorescence from 0 to 4 h was evaluated for each aaRS activity. The concentration of each aaRS was plotted against aaRS activity, and the specific activity was calculated using linear regression.

### Serial-dilution experiment

In round 1, a single or multiple DNA encoding each aaRS (aaRS-DNA) and firefly luciferase (Fluc-DNA) were incubated in the customized PURE system (version 2) containing all 20 aaRS proteins (the complete composition is shown in Table S2) at 30 °C for 8 h. The total DNA concentrations were 5 nM (Fig. 4), 2 nM (Fig. 7), 2-7.5 nM (Fig. S8), and 2-3 nM (Fig. S9). The concentration ratio of each aaRS-DNA is presented in Tables S3 and S4. In the subsequent rounds, the reaction solution after incubation in the previous round was diluted 2-, 2.5-, (Figs. 7-8 and S8-9), or 5-(Fig. 4) fold with another customized PURE system (version 2, PUREΔaaRS) that lacks aaRSs of interest and contains aaRS-DNA and Fluc-DNA at the same concentrations as round 1, and incubated again at 30 °C for 8 h. The serial dilution process was repeated during the indicated rounds. At the end of each round, 1 µL of an aliquot of the reaction mixture was added to 30 µL of the Luciferase Assay Reagent (Promega), and luminescence was measured as translational activity using a luminometer (GloMax, Promega). For negative control experiments, DNA encoding GFP (GFP-DNA) was used instead of aaRS-DNA. Additionally, as a positive control, we conducted the same serial dilution experiment using GFP-DNA and a customized PURE system (version 2) containing 20 aaRS proteins (Full PURE in Fig. 4).

### An assay of the inhibitory effect of aaRS DNA on translation (Fig. 5)

To examine the inhibitory effect (Fig. 5A), aaRS-DNA (5 nM) and Fluc-DNA (0.01 nM) were incubated at 30°C for 8 h in a customized PURE system (version 2) containing all aaRS proteins. For the assay in Figs. 5B and 5C, aaRS-DNA (1 nM), GFP-DNA (4 nM), and Fluc-DNA (0.01 nM) were incubated in the PURE system (version 2). As controls, GFP-DNA (5 nM) and Fluc-DNA (0.01 nM) were incubated in the PURE system (version 2). EF-P (1 µM, GeneFrontier) was added in Fig. 5C. After incubation, 1 µL of an aliquot of the reaction mixture was added to 30 µL of the luciferase assay reagent (Promega), and luminescence was measured as translational activity using a luminometer (GloMax, Promega).

### Determination of DNA concentration ratio

The concentration ratio of each aaRS in the aaRS-DNA mix used in serial dilution experiments was determined as follows. First, we defined a requirement index, R, which reflects the requirement of each aaRS-DNA for regeneration. The requirement index should be proportional to each concentration and the specific activity of the endogenous aaRS included in the PURE system but inversely proportional to each concentration and the specific activity of the aaRS expressed in the PURE system. Therefore, R can be defined using the H/L ratio (concentration ratio of expressed to endogenous aaRS) and specific activity (*A_exp_/A_end_*) obtained in Fig. 1 as follows (Equation 2):

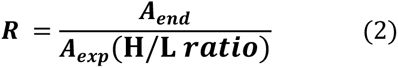

The concentration of each DNA was determined as follows (Equation 3). The total DNA concentration differed depending on the experimental conditions used.

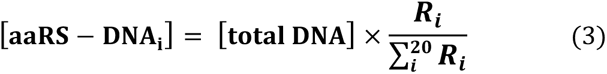

In Figs. 6C, 6D, and 7, aaRS-DNA concentrations were calibrated by multiplying the calibration value (*C_DNA_*) as follows (Equation 4):

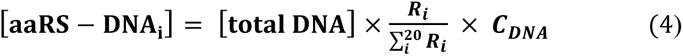

The calibration value, which represents the insufficiency of each aaRS-DNA, is equal to the average of the y-axis values in Fig. 6B (i.e., the dotted line) divided by each y-axis value. After each DNA concentration ([aaRS-DNA_i_]) was calculated, the DNA concentrations were normalized to ensure that the sum of the calculated DNA concentrations was equal to the target total DNA concentration.

### The prediction of the translation amount in co-expressing 20 aaRSs used in Figs. 6B and 6D

The translation amount was predicted as follows based on the translation efficiency (T) measured in Fig. 1B, the DNA concentration ratio, and the activity improvement rate of the PURE system obtained in Fig. 2C (14.4) (Equation 5).

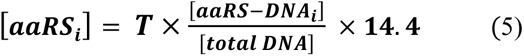

## Supplementary Materials

Supplementary material for this article is available at XXX

## Acknowledgements

We thank Mrs. Kayo Aoyama and Ayu Saito for technical support.

## Author contributions

K.H., K.K., N.I., and Y.S. designed the study and wrote the manuscript. K. H., K. M., N.I. performed experiments and analyses.

## Data and materials availability

Data is available on request from the authors.

## References

1. J. W. Szostak, D. P. Bartel, P. L. Luisi, Synthesizing life. Nature 409, 387–390 (2001).

2. A. C. Forster, G. M. Church, Towards synthesis of a minimal cell. Mol Syst Biol 2 (2006).

3. N. Ichihashi, What can we learn from the construction of in vitro replication systems? Ann N Y Acad Sci 1447, 144–156 (2019).

4. M. Forlin, R. Lentini, S. S. Mansy, Cellular imitations. Curr Opin Chem Biol 16, 586–592 (2012).

5. B. C. Buddingh’, J. C. M. van Hest, Artificial Cells: Synthetic Compartments with Life-like Functionality and Adaptivity. Acc Chem Res 50, 769–777 (2017).

6. N. A. Yewdall, A. F. Mason, J. C. M. Van Hest, The hallmarks of living systems: Towards creating artificial cells. Interface Focus 8, 20180023 (2018).

7. A. D. Silverman, A. S. Karim, M. C. Jewett, Cell-free gene expression: an expanded repertoire of applications. Nat Rev Genet 21, 151–170 (2020).

8. V. Noireaux, A. P. Liu, The New Age of Cell-Free Biology. Annu Rev Biomed Eng 22, 51–77 (2020).

9. L. Damiano, P. Stano, On the “Life-Likeness” of Synthetic Cells. Front Bioeng Biotechnol 8, 953 (2020).

10. E. Cho, Y. Lu, Compartmentalizing Cell-Free Systems: Toward Creating Life-Like Artificial Cells and Beyond. ACS Synth Biol 9, 2881–2901 (2020).

11. I. Ivanov, S. L. Castellanos, S. Balasbas, L. Otrin, N. Maruscaroniccaron, T. Vidakovicacute-Koch, K. Sundmacher, Bottom-Up Synthesis of Artificial Cells: Recent Highlights and Future Challenges. Annu Rev Chem Biomol Eng 12, 287–308 (2021).

12. C. Wang, J. Yang, Y. Lu, Modularize and Unite: Toward Creating a Functional Artificial Cell. Front Mol Biosci 8, 781986 (2021).

13. Y. Lyu, R. Peng, H. Liu, H. Kuai, L. Mo, D. Han, J. Li, W. Tan, Protocells programmed through artificial reaction networks. Chem Sci 11, 631–642 (2020).

14. L. Olivi, M. Berger, R. N. P. Creyghton, N. De Franceschi, C. Dekker, B. M. Mulder, N. J. Claassens, P. Rein ten Wolde, J. van der Oost, Towards a synthetic cell cycle. Nat Commun 12, 4531 (2015).

15. J. C. Blain, J. W. Szostak, Progress toward synthetic cells. Annu Rev Biochem 83, 615–640 (2014).

16. N. J. Gaut, K. P. Adamala, Reconstituting Natural Cell Elements in Synthetic Cells. Adv Biol 5, 1–20 (2021).

17. J. De Capitani, H. Mutschler, The Long Road to a Synthetic Self-Replicating Central Dogma. Biochemistry 62, 1221–1232 (2023).

18. M. Su’etsugu, H. Takada, T. Katayama, H. Tsujimoto, Exponential propagation of large circular DNA by reconstitution of a chromosome-replication cycle. Nucleic Acids Res 45, 11525–11534 (2017).

19. P. van Nies, I. Westerlaken, D. Blanken, M. Salas, M. Mencía, C. Danelon, Self-replication of DNA by its encoded proteins in liposome-based synthetic cells. Nat Commun 9, 1583 (2018).

20. Y. Sakatani, T. Yomo, N. Ichihashi, Self-replication of circular DNA by a self-encoded DNA polymerase through rolling-circle replication and recombination. Sci Rep 8, 13089 (2018).

21. H. Okauchi, Y. Sakatani, K. Otsuka, N. Ichihashi, Minimization of Elements for Isothermal DNA Replication by an Evolutionary Approach. ACS Synth Biol 9, 1771–1780 (2020).

22. S. Berhanu, T. Ueda, Y. Kuruma, Artificial photosynthetic cell producing energy for protein synthesis. Nat Commun 10, 1325 (2019).

23. K. Y. Lee, S.-J. Park, K. A. Lee, S.-H. Kim, H. Kim, Y. Meroz, L. Mahadevan, K.-H. Jung, T. K. Ahn, K. K. Parker, K. Shin, Photosynthetic artificial organelles sustain and control ATP-dependent reactions in a protocellular system. Nat Biotechnol 36, 530–535 (2018).

24. S. Giaveri, N. Bohra, C. Diehl, H. Y. Yang, M. Ballinger, N. Paczia, T. Glatter, T. J. Erb, Integrated translation and metabolism in a partially self-synthesizing biochemical network. Science(1979) 385, 174–178 (2024).

25. S. Sugii, K. Hagino, R. Mizuuchi, N. Ichihashi, Cell-free expression of RuBisCO for ATP production in the synthetic cells. Synth Biol 8, ysad016 (2023).

26. D. Blanken, D. Foschepoth, A. C. Serrão, C. Danelon, Genetically controlled membrane synthesis in liposomes. Nat Commun 11, 4317 (2020).

27. S. Eto, R. Matsumura, Y. Shimane, M. Fujimi, S. Berhanu, T. Kasama, Y. Kuruma, Phospholipid synthesis inside phospholipid membrane vesicles. Commun Biol 5, 1016 (2022).

28. S. Kohyama, A. Merino-Salomón, P. Schwille, In vitro assembly, positioning and contraction of a division ring in minimal cells. Nat Commun 13, 6098 (2022).

29. G. Sato, S. Miyazawa, N. Doi, K. Fujiwara, Cell-Free Protein Expression by a Reconstituted Transcription–Translation System Energized by Sugar Catabolism. Molecules 29 (2024).

30. E. Karzbrun, A. M. Tayar, V. Noireaux, R. H. Bar-Ziv, Programmable on-chip DNA compartments as artificial cells. Science (1979) 345, 829–832 (2014).

31. M. A. Boyd, W. Thavarajah, J. B. Lucks, N. P. Kamat, Robust and tunable performance of a cell-free biosensor encapsulated in lipid vesicles. Sci Adv 9, eadd6605 (2024).

32. Z.-H. Luo, C. Chen, Q.-H. Zhao, N.-N. Deng, Functional metal-phenolic cortical cytoskeleton for artificial cells. Sci Adv 10, eadj4047 (2024).

33. S. Takada, N. Yoshinaga, N. Doi, K. Fujiwara, Mode selection mechanism in traveling and standing waves revealed by Min wave reconstituted in artificial cells. Sci Adv 8, eabm8460 (2024).

34. M. Levy, R. Falkovich, S. S. Daube, R. H. Bar-Ziv, Autonomous synthesis and assembly of a ribosomal subunit on a chip. Sci Adv 6, eaaz6020 (2024).

35. Y. Shimizu, A. Inoue, Y. Tomari, T. Suzuki, T. Yokogawa, K. Nishikawa, T. Ueda, Cell-free translation reconstituted with purified components. Nat Biotechnol 19, 751–755 (2001).

36. T. Awai, N. Ichihashi, T. Yomo, Activities of 20 aminoacyl-tRNA synthetases expressed in a reconstituted translation system in Escherichia coli. Biochem Biophys Rep 3, 140–143 (2015).

37. E. Wei, D. Endy, Experimental tests of functional molecular regeneration via a standard framework for coordinating synthetic cell building. bioRxiv, 2021.03.03.433818 (2021).

38. J. Li, W. Haas, K. Jackson, E. Kuru, M. C. Jewett, Z. H. Fan, S. Gygi, G. M. Church, Cogenerating Synthetic Parts toward a Self-Replicating System. ACS Synth Biol 6, 1327–1336 (2017).

39. K. Libicher, R. Hornberger, M. Heymann, H. Mutschler, In vitro self-replication and multicistronic expression of large synthetic genomes. Nat Commun 11, 1–8 (2020).

40. A. Doerr, D. Foschepoth, A. C. Forster, C. Danelon, In vitro synthesis of 32 translation-factor proteins from a single template reveals impaired ribosomal processivity. Sci Rep 11, 1–12 (2021).

41. K. Libicher, H. Mutschler, Probing self-regeneration of essential protein factors required for: In vitro translation activity by serial transfer. Chemical Communications 56, 15426–15429 (2020).

42. B. Lavickova, N. Laohakunakorn, S. J. Maerkl, A partially self-regenerating synthetic cell. Nat Commun 11, 6340 (2020).

43. K. Hagino, N. Ichihashi, In Vitro Transcription/Translation-Coupled DNA Replication through Partial Regeneration of 20 Aminoacyl-tRNA Synthetases. ACS Synth Biol 12, 1252–1263 (2023).

44. Y. Shimizu, T. Kanamori, T. Ueda, Protein synthesis by pure translation systems. Methods 36, 299–304 (2005).

45. P. O. Olins, C. S. Devine, S. H. Rangwala, K. S. Kavka, The T7 phage gene 10 leader RNA, a ribosome-binding site that dramatically enhances the expression of foreign genes in Escherichia coli. Gene 73, 227–235 (1988).

46. T. Fuse-Murakami, R. Matsumoto, T. Kanamori, N-Terminal Amino Acid Affects the Translation Efficiency at Lower Temperatures in a Reconstituted Protein Synthesis System. Int J Mol Sci 25 (2024).

47. C.-M. Zhang, T. Christian, K. J. Newberry, J. J. Perona, Y.-M. Hou, Zinc-mediated Amino Acid Discrimination in Cysteinyl-tRNA Synthetase. J Mol Biol 327, 911–917 (2003).

48. J. Liu, Y. Gagnon, J. Gauthier, L. Furenlid, P.-J. L’Heureux, M. Auger, O. Nureki, S. Yokoyama, J. Lapointe, The Zinc-binding Site of Escherichia coli Glutamyl-tRNA Synthetase Is Located in the Acceptor-binding Domain. Journal of Biological Chemistry 270, 15162–15169 (1995).

49. Y. Kazuta, T. Matsuura, N. Ichihashi, T. Yomo, Synthesis of milligram quantities of proteins using a reconstituted in vitro protein synthesis system. J Biosci Bioeng 118, 554–557 (2014).

50. J. S. Madin, D. A. Nielsen, M. Brbic, R. Corkrey, D. Danko, K. Edwards, M. K. M. Engqvist, N. Fierer, J. L. Geoghegan, M. Gillings, N. C. Kyrpides, E. Litchman, C. E. Mason, L. Moore, S. L. Nielsen, I. T. Paulsen, N. D. Price, T. B. K. Reddy, M. A. Richards, E. P. C. Rocha, T. M. Schmidt, H. Shaaban, M. Shukla, F. Supek, S. G. Tetu, S. Vieira-Silva, A. R. Wattam, D. A. Westfall, M. Westoby, A synthesis of bacterial and archaeal phenotypic trait data. Sci Data 7, 170 (2020).

51. R. Krafczyk, F. Qi, A. Sieber, J. Mehler, K. Jung, D. Frishman, J. Lassak, Proline codon pair selection determines ribosome pausing strength and translation efficiency in bacteria. Commun Biol 4, 589 (2021).

52. K. R. Hummels, D. B. Kearns, Translation elongation factor P (EF-P). FEMS Microbiol Rev 44, 208–218 (2020).

53. S. Ude, J. Lassak, A. L. Starosta, T. Kraxenberger, D. N. Wilson, K. Jung, Translation Elongation Factor EF-P Alleviates Ribosome Stalling at Polyproline Stretches. Science (1979) 339, 82–85 (2013).

54. M. C. J. Yip, S. Shao, Detecting and Rescuing Stalled Ribosomes. Trends Biochem Sci 46, 731–743 (2021).

55. T. Inada, R. Beckmann, Mechanisms of Translation-coupled Quality Control. J Mol Biol 436, 168496 (2024).

56. R. B. Ganesh, S. J. Maerkl, Towards Self-regeneration: Exploring the Limits of Protein Synthesis in the Protein Synthesis Using Recombinant Elements (PURE) Cell-free Transcription–Translation System. ACS Synth Biol 13, 2555–2566 (2024).

57. K. Seo, N. Ichihashi, Investigation of Compatibility between DNA Replication, Transcription, and Translation for in Vitro Central Dogma. ACS Synth Biol 12, 1813–1822 (2023).

58. Y. Ito, T. Kawama, I. Urabe, T. Yomo, Evolution of an Arbitrary Sequence in Solubility. J Mol Evol 58, 196–202 (2004).

59. Y. Murase, H. Nakanishi, G. Tsuji, T. Sunami, N. Ichihashi, In Vitro Evolution of Unmodified 16S rRNA for Simple Ribosome Reconstitution. ACS Synth Biol 7, 576–583 (2018).

60. K. Takai, CodHonEditor: Spreadsheets for Codon Optimization and Editing of Protein Coding Sequences. Nucleosides Nucleotides Nucleic Acids 35, 223–232 (2016).

61. S. Fujii, T. Matsuura, T. Sunami, T. Nishikawa, Y. Kazuta, T. Yomo, Liposome display for in vitro selection and evolution of membrane proteins. Nat Protoc 9, 1578–1591 (2014).

62. T. Masuda, M. Tomita, Y. Ishihama, Phase Transfer Surfactant-Aided Trypsin Digestion for Membrane Proteome Analysis. J Proteome Res 7, 731–740 (2008).

63. J. Adachi, K. Hashiguchi, M. Nagano, M. Sato, A. Sato, K. Fukamizu, Y. Ishihama, T. Tomonaga, Improved Proteome and Phosphoproteome Analysis on a Cation Exchanger by a Combined Acid and Salt Gradient. Anal Chem 88, 7899–7903 (2016).

