## Supplemental information 1 for "Sustainable Regeneration of 20 Aminoacyl-tRNA Synthetases in a Reconstituted System Toward Self-Synthesizing Artificial Systems"

**This file includes:**

Fig. S1-S9

Table. S1-S5

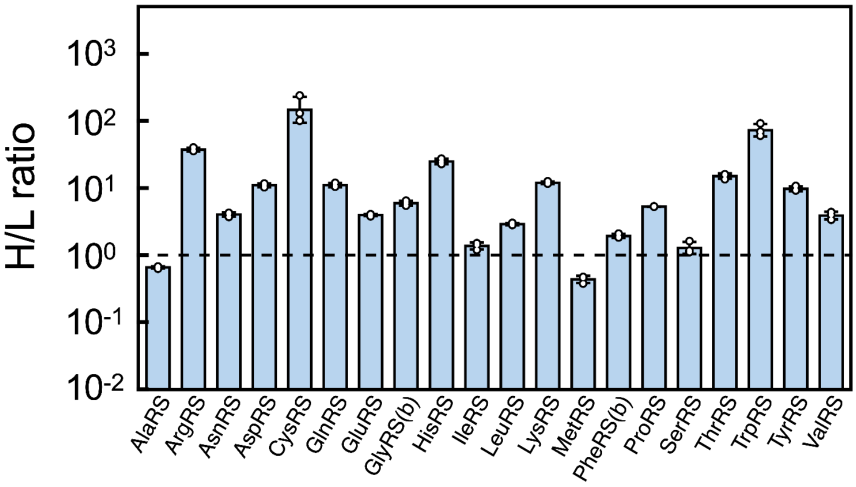


**Fig. S1. H/L ratio of Fig. 1B**

Each point represents an individual detected peptide, with error bars indicating standard deviation.

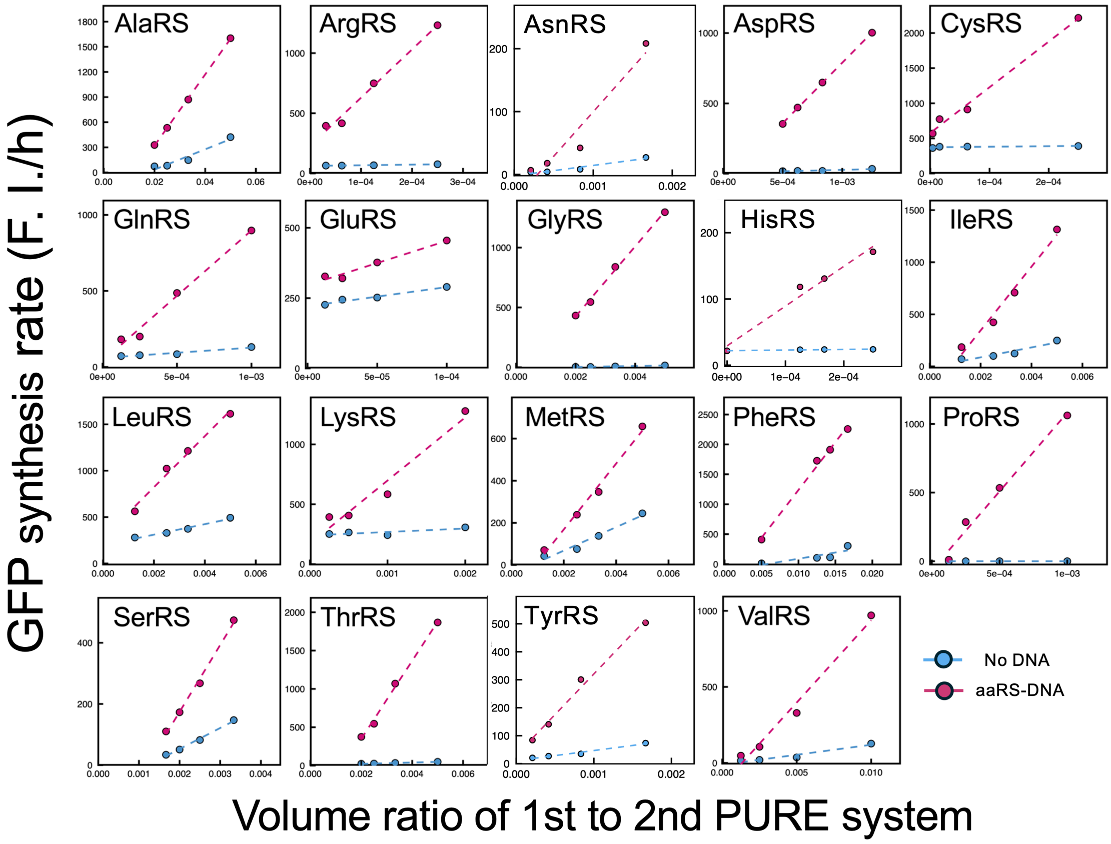


**Fig. S2. Raw data plot of activity assay of Fig. 1C**

The y-axis shows the GFP synthesis rate and the x-axis shows the volume ratio of the 1st to 2$^{nd}$ PURE system after dilution. The slope was used as $k_{exp+end}$ (aaRS-DNA) and $k_{end}$ (No DNA).

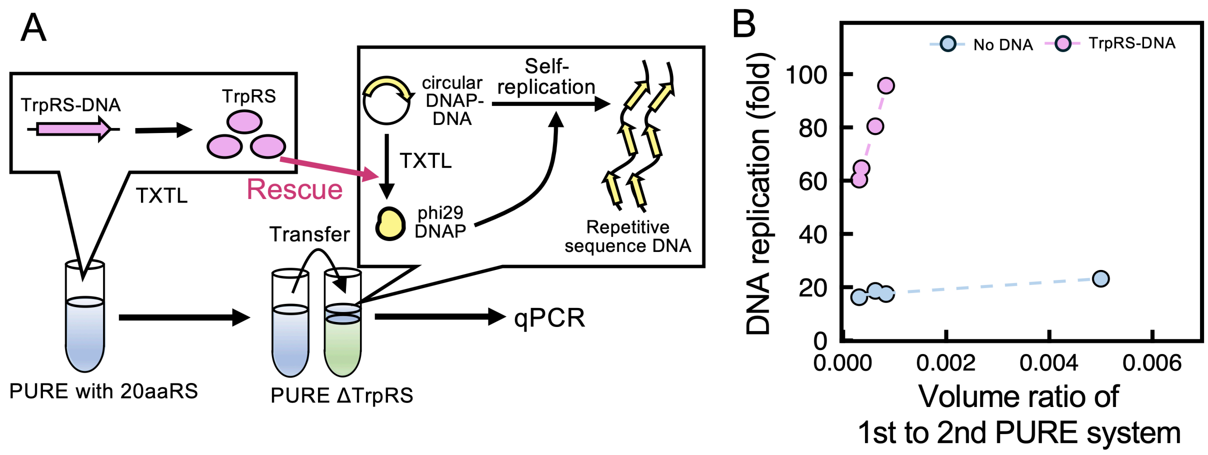


**Fig. S3. TrpRS activity assay using self-replicating DNA**

(A) Scheme of the assay. The PURE system (version 1), in which TrpRS-DNA (5 nM) was incubated at 30°C for 8 h, was diluted with a dilution buffer (50 mM HEPES-KOH (pH7.6), 100 mM KCl, 10 mM $MgCl_2$, 7 mM 2-mercaptoethanol, 5 mg/mL BSA, 10 mM DTT, and 30% glycerol), added to the next PURE system lacking TrpRS and containing a circular phi29 DNAP-DNA that encodes phi29 DNA polymerase, and incubated again at 30°C for 16 h. After incubation, the replication rate of phi29 DNAP-DNA was quantified by quantitative PCR using the primers 5 (AGGGTATGGGCGTATGGTTATATG) and 6 (TGTCCCATGCGAGATATGATCG). (B) Raw data plots of the TrpRS activity assay. The y-axis shows the fold of DNA replication and the x-axis shows the volume ratio of the 1st to 2nd PURE system after dilution.

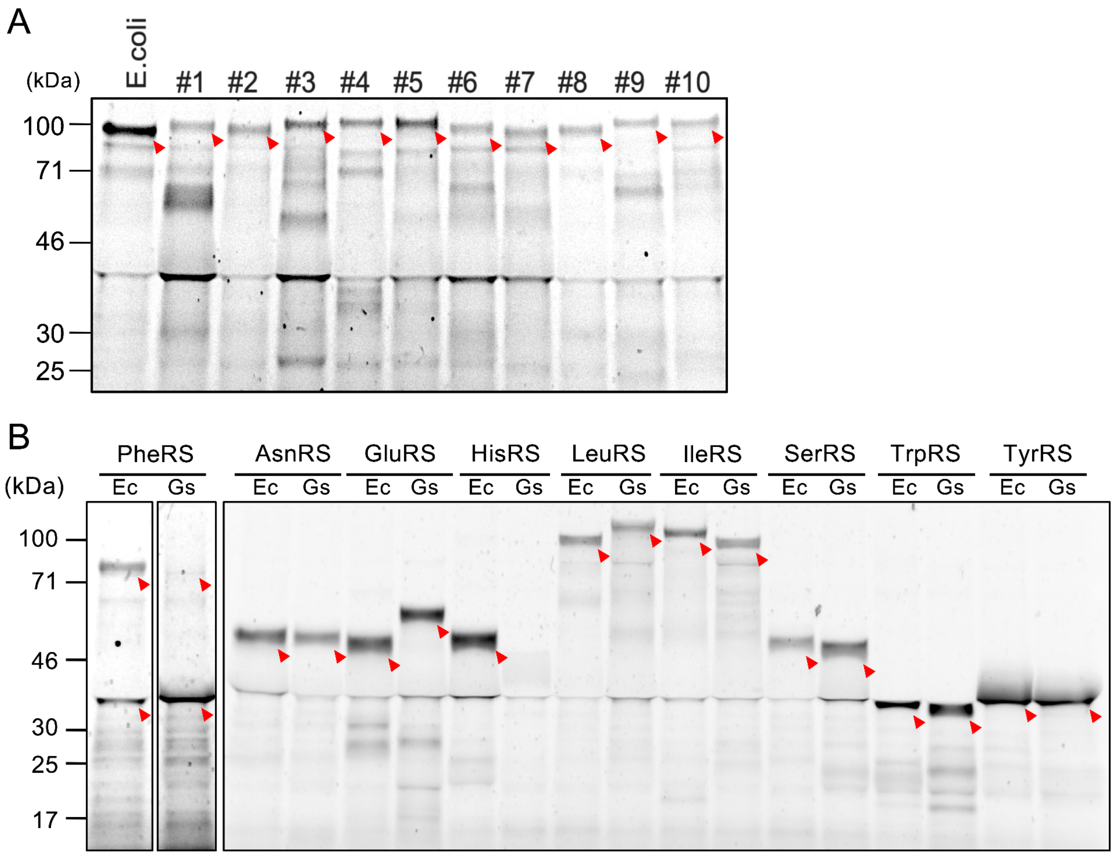


**Fig. S4. Expression of aaRSs derived from other bacteria in the PURE system**

(A) A gel image of SDS-PAGE for AlaRSs derived from ten bacteria (Fig. 3A). (B) Gel images of SDS-PAGE of nine aaRSs derived from *E. coli* (Ec) and *G. stearothermophilus* (Gs). The expected bands for each aaRS protein are indicated by the red arrowheads.

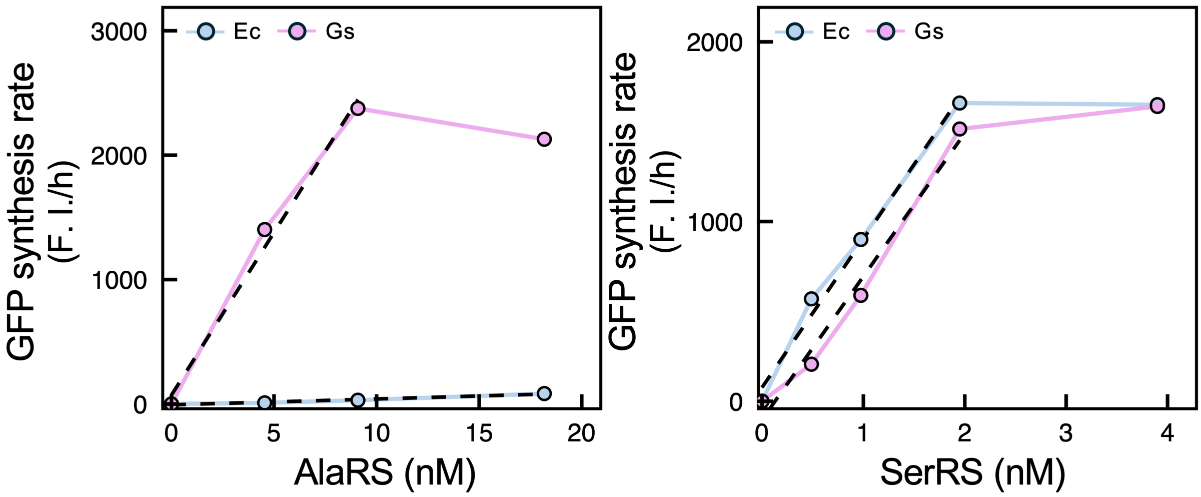


**Fig. S5. Raw data plots of Fig. 3F.**

GFP synthesis rates were measured by the same method as shown in Fig. 1A(2) by using purified Ec-and Gs-AlaRSs or SerRSs instead of expressing in the 1st PURE system. The X-axis shows the aaRS concentration, and the Y-axis shows the rate of increase in GFP fluorescence. The dotted line indicates the regression line used to calculate the specific activity.

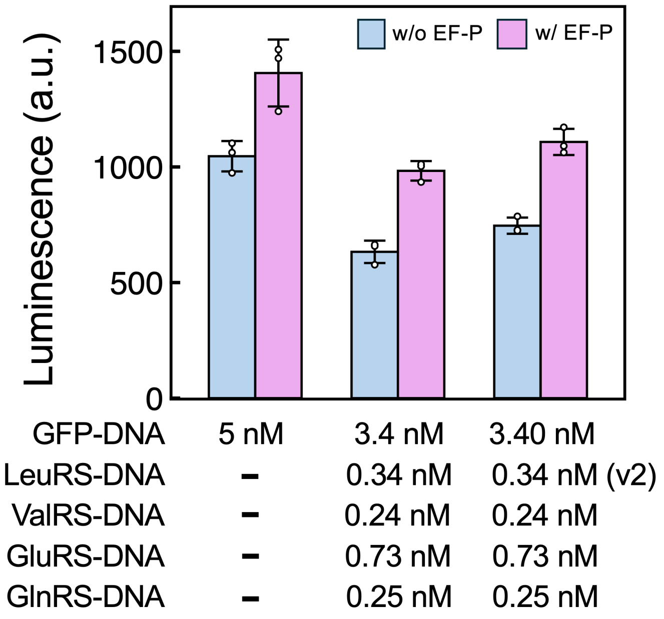


**Fig. S6. Total inhibitory effect by four aaRS**

Translation inhibition by four inhibitory aaRS was measured at realistic concentrations using the same method as in Fig. 5. Each aaRS-DNA concentration was based on the DNA concentration ratio for 20 aaRS regenerations before calibration (Table S4, total DNA 5 nM).

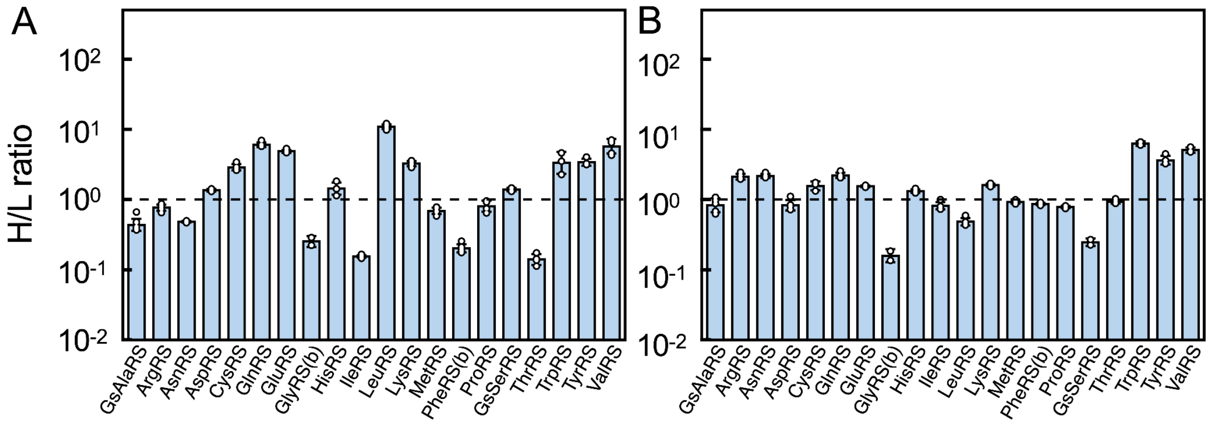


**Fig. S7. The H/L ratios are shown in Figure 6.**

Each point represents an individual detected peptide, with error bars showing the standard deviation before (A) and after (B) calibration of aaRS-DNA concentrations.

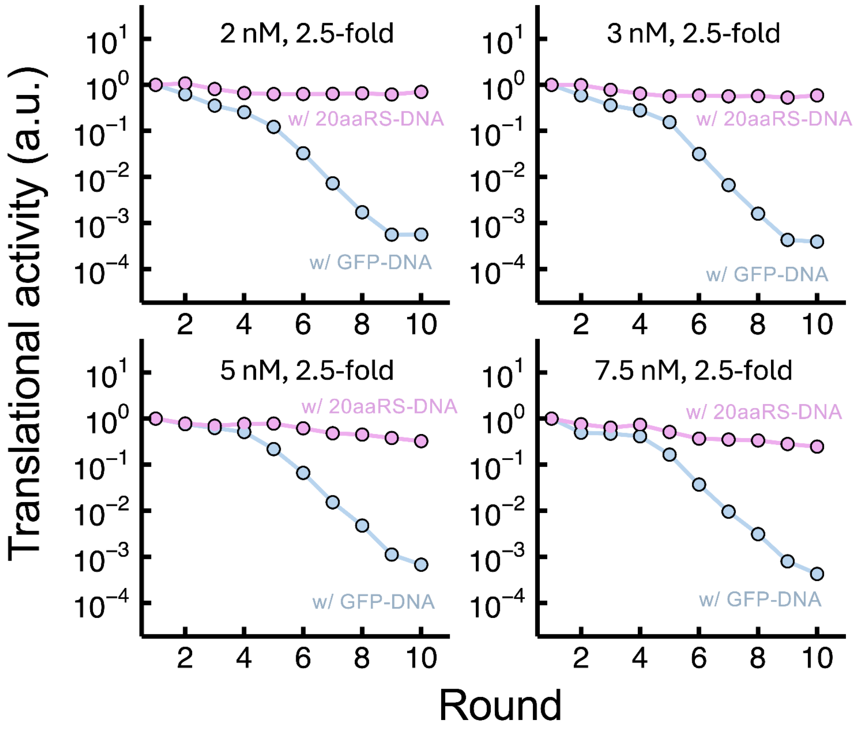


**Fig. S8. Effect of total DNA concentration and dilution rates on 20 aaRS regeneration**.

Experiments were performed using the method shown in Fig. 7A.

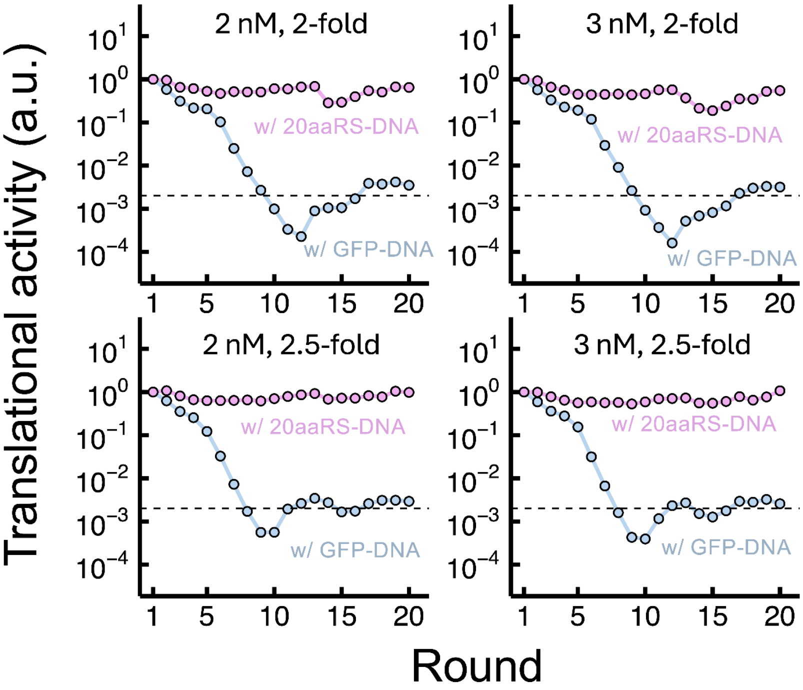


**Fig. S9. Sustainable regeneration of 20 aaRSs under different conditions.**

The serial dilution experiments shown in Fig. 7A were performed under different conditions (3 nM aaRS-DNA and/or 2-fold dilution). In either case, translation activity was maintained at a certain level until 20 rounds. For comparison, the same data as Fig. 7B (2 nM aaRS-DNAs, 2.5-fold dilution) is shown

**Table S1. Sequences property and translation efficiency of 20 aaRS**

| aaRS | Translation (nM) | 5’UTR | GC in 7aa (%) | P repeat |
|---|---|---|---|---|
| CysRS | 3750.36 | Long | 23.18 | - |
| HisRS | 2127.40 | Long | 38.18 | - |
| TrpRS | 2048.37 | Long | 38.1 | PP |
| AsnRS | 1693.34 | Long | 33.3 | - |
| TyrRS | 1481.89 | Long | 38.1 | - |
| LysRS | 1375.02 | Long | 42.9 → 33.3 | PP |
| AspRS | 1338.06 | Long | 38.1 | PP |
| ThrRS | 1262.79 | Long | 33.3 | - |
| ArgRS | 1154.39 | Long (mutation) | 42.9 → 33.3 | - |
| GluRS | 917.14 | Long | 33.3 | PP |
| ProRS | 879.60 | Long | 47.6 → 28.6 | - |
| GlnRS | 662.32 | Long | 61.9 → 42.9 | PP |
| GlyRS(a) | 514.55 | Long | 38.1 | - |
| IleRS | 500.32 | Long | 38.1 | PP |
| AlaRS | 476.60 | Long | 33.3 | PP |
| PheRS(a) | 259.28 | Long | 33.3 | PP, PP |
| LeuRS | 119.00 | Short | 47.6 →42.9 | - |
| SerRS | 100.70 | Short | 42.9 →33.3 | - |
| ValRS | 67.09 | Short | 33.3 | PPP |
| MetRS | 47.64 | Short | 39 | - |

**Table S2. Composition of the customized PURE system versions 1 and 2**

| | | | |
|---|---|---|---|
| Initiation factor 1 | 25 µM | ValRS | 17 nM |
| Initiation factor 2 | 1 µM | Methionyl-tRNA formyltransferase | 590 nM |
| Initiation factor 3 | 4.9 µM | Myokinase | 1.4 µM |
| Elongation factor G | 1.1 µM | Creatine kinase | 250 nM |
| Elongation factor Tu | 80 µM | Nucleoside diphosphate kinase | 16 nM |
| Elongation factor Ts | 3.3 µM | Pyrophosphatase | 41 nM |
| Release factor 1 | 49 nM | Trigger factor | 1 µM |
| Release factor 2 | 48 nM | *E. coli* DEAH type RNA helicase A | 100 nM |
| Release factor 3 | 170 nM | 70S ribosome | 1 µM |
| Ribosome recycling factor | 3.9 nM | | |
| AlaRS | 730 nM | Tyrosine | 0.3 mM |
| ArgRS | 31 nM | Cysteine | 0.3 mM |
| AsnRS | 420 nM | 18 other amino acids | 0.36 mM |
| AspRS | 120 nM | tRNA mix (Roche, *E. coli*) | 0.52 (3.12) mg/mL |
| CysRS | 24 nM | ATP | 0.375 (3.75) mM |
| GlnRS | 60 nM | GTP | 0.25 (2.5) mM |
| GluRS | 230 nM | CTP | 0.125 (1.25) mM |
| GlyRS | 86 nM | UTP | 0.125 (1.25) mM |
| HisRS | 85 nM | N-2-hydroxyethylpiperazine-N'-2-ethanesulfonic acid (pH 7.6) | 100 mM |
| IleRS | 370 nM | Glutamate acid potassium salt | 70 (280) mM |
| LeuRS | 41 nM | Spermidine | 0.375 (1.5) mM |
| LysRS | 120 nM | Magnesium acetate | 10.7 mM |
| MetRS | 110 nM | Creatine phosphate | 25 mM |
| PheRS | 130 nM | Dithiothreitol | 6 mM |
| ProRS | 170 nM | 10-formyl-5,6,7,8-tetrahydro folic acid | 10 µg/mL |
| SerRS | 78 nM | Yeast inorganic pyrophosphatase (NEB) | 0.2 mU/µL |
| ThrRS | 84 nM | RNase inhibitor (Promega) | 0.1 U/µL |
| TrpRS | 28 nM | T7 RNA polymerase (Takara) | 0.42 (1.77) U/µL |
| TyrRS | 150 nM | Zinc acetate | 0 (1) mM |

The concentrations in version 2 are shown in parentheses.

**Table S3. DNA concentration ratio in Fig. 4**

| aaRS | DNA (nM) | | | | | | |
|---|---|---|---|---|---|---|---|
| | Control | 1 aaRS | 5 aaRS | 10 aaRS | | | |
| | | | | Group1 | Group2 | Group3 | Group4 |
| GFP | 5.00 | - | - | - | - | - | - |
| GsAlaRS | - | - | - | - | 0.46 | - | 0.10 |
| ArgRS | - | - | - | - | 0.29 | 0.07 | - |
| AsnRS | - | - | 0.83 | 0.59 | - | 0.82 | 0.71 |
| AspRS | - | - | - | - | 0.64 | - | 0.14 |
| CysRS | - | - | - | - | 0.14 | 0.03 | - |
| GlnRS | - | - | - | 0.30 | - | - | - |
| GluRS | - | - | 1.21 | 0.87 | - | 3.74 | 1.04 |
| GlyRS | - | - | - | - | 0.38 | 0.09 | - |
| HisRS | - | - | - | - | 0.39 | 0.09 | - |
| IleRS | - | 5.00 | 1.14 | 0.82 | - | 2.15 | 0.98 |
| LeuRS | - | - | - | 0.41 | - | - | - |
| LysRS | - | - | - | - | 0.83 | - | 0.18 |
| MetRS | - | - | - | - | 0.93 | - | 0.20 |
| PheRS | - | - | 0.93 | 0.66 | - | 1.33 | 0.80 |
| ProRS | - | - | - | 0.22 | - | - | - |
| GsSerRS | - | - | - | 0.21 | - | - | - |
| ThrRS | - | - | - | - | 0.44 | 0.11 | - |
| TrpRS | - | - | - | - | 0.50 | - | 0.11 |
| TyrRS | - | - | 0.89 | 0.63 | - | 1.04 | 0.76 |
| ValRS | - | - | - | 0.29 | - | - | - |
| Luciferase | 0.01 | 0.01 | 0.01 | 0.01 | 0.01 | 0.01 | 0.01 |

**Table S4. DNA concentration ratio for 20 aaRS regeneration**

| aaRS | DNA (nM) | |
|---|---|---|
| | Before | After calibration |
| GsAlaRS | 0.07 | 0.01 |
| ArgRS | 0.04 | 0.02 |
| AsnRS | 0.5 | 0.33 |
| AspRS | 0.1 | 0.01 |
| CysRS | 0.02 | 0.004 |
| GlnRS | 0.26 | 0.02 |
| GluRS | 0.73 | 0.07 |
| GlyRS | 0.06 | 0.01 |
| HisRS | 0.06 | 0.01 |
| IleRS | 0.69 | 0.68 |
| LeuRS | 0.35 | 0.01 (v2) |
| LysRS | 0.13 | 0.02 |
| MetRS | 0.14 | 0.03 |
| PheRS | 0.56 | 0.48 |
| ProRS | 0.19 | 0.05 |
| GsSerRS | 0.17 | 0.01 |
| ThrRS | 0.07 | 0.07 |
| TrpRS | 0.08 | 0.02 |
| TyrRS | 0.54 | 0.13 |
| ValRS | 0.24 | 0.01 |

### Table S5. Composition of the customized PURE system used in Fig. 7

| | | | |
|---|---|---|---|
| Initiation factor 1 | 25 µM | ValRS | 17 nM |
| Initiation factor 2 | 1 µM | Methionyl-tRNA formyltransferase | 590 nM |
| Initiation factor 3 | 4.9 µM | Myokinase | 1.4 µM |
| Elongation factor G | 1.1 µM | Creatine kinase | 250 nM |
| Elongation factor Tu | 80 µM | Nucleoside diphosphate kinase | 16 nM |
| Elongation factor Ts | 3.3 µM | Pyrophosphatase | 41 nM |
| Release factor 1 | 49 nM | Trigger factor | 1 µM |
| Release factor 2 | 48 nM | *E. coli* DEAH type RNA helicase A | 100 nM |
| Release factor 3 | 170 nM | 70S ribosome | 1 µM |
| Ribosome recycling factor | 3.9 nM | Elongation factor P | 1 µM |
| AlaRS | 730 nM | Tyrosine | 0.3 mM |
| ArgRS | 31 nM | Cysteine | 0.3 mM |
| AsnRS | 420 nM | 18 other amino acids | 0.36 mM |
| AspRS | 120 nM | tRNA mix (Roche, *E. coli*) | 3.12 mg/mL |
| CysRS | 24 nM | ATP | 3.75 mM |
| GlnRS | 60 nM | GTP | 2.5 mM |
| GluRS | 230 nM | CTP | 1.25 mM |
| GlyRS | 86 nM | UTP | 1.25 mM |
| HisRS | 85 nM | N-2-hydroxyethylpiperazine-N'-2-ethanesulfonic acid (pH 7.6) | 100 mM |
| IleRS | 370 nM | Glutamate acid potassium salt | 280 mM |
| LeuRS | 41 nM | Spermidine | 1.5 mM |
| LysRS | 120 nM | Magnesium acetate | 14.7 mM |
| MetRS | 110 nM | Zinc acetate | 1 mM |
| PheRS | 130 nM | Creatine phosphate | 25 mM |
| ProRS | 170 nM | Dithiothreitol | 6 mM |
| SerRS | 78 nM | 10-formyl-5,6,7,8-tetrahydro folic acid | 10 µg/mL |
| ThrRS | 84 nM | RNase inhibitor (Promega) | 0.1 U/µL |
| TrpRS | 28 nM | T7 RNA polymerase (Takara) | 1.77 U/µL |
| TyrRS | 150 nM | | |
